# Co-ordinated regulation of cell survival and cell cycle pathways by DDR2-dependent SRF transcription factor in cardiac fibroblasts

**DOI:** 10.1101/857037

**Authors:** Allen Sam Titus, V Harikrishnan, Shivakumar Kailasam

**Affiliations:** Division of Cellular and Molecular Cardiology, Sree Chitra Tirunal Institute for Medical Sciences and Technology, Trivandrum, India

**Keywords:** Cardiac fibroblasts, Discoidin Domain Receptor 2 (DDR2), ERK1/2 MAPK, Serum Response Factor (SRF), cIAP2, FoxO3a, Skp2, p27, apoptosis resistance, G1-S transition

## Abstract

Relative resistance to apoptosis and the ability to proliferate and produce a collagen-rich scar determine the critical role of cardiac fibroblasts in wound healing and tissue remodeling following myocardial injury. Identification of cardiac fibroblast-specific factors and mechanisms underlying these aspects of cardiac fibroblast function is therefore of considerable scientific and clinical interest. In the present study, gene knockdown and over-expression approaches, and promoter binding assays, showed that DDR2, a mesenchymal cell-specific collagen receptor tyrosine kinase localized predominantly in fibroblasts in the heart, acts via ERK1/2 MAPK-activated SRF transcription factor to enhance the expression of anti-apoptotic cIAP2 in cardiac fibroblasts, conferring resistance against oxidative injury.

Further, DDR2 was found to act via ERK1/2 MAPK-activated SRF to transcriptionally up regulate Skp2 that in turn facilitated post-translational degradation of p27, the cyclin dependent kinase inhibitor that causes cell cycle arrest, to promote G1-S transition, as evidenced by Rb phosphorylation, increased PCNA levels and flow cytometry. DDR2-dependent ERK1/2 MAPK activation also suppressed FoxO3a-mediated transcriptional induction of p27. Inhibition of the binding of collagen type I to DDR2 using WRG-28 indicated the obligate role of collagen type I in the activation of DDR2 and its regulatory role in cell survival and cell cycle protein expression. Notably, DDR2 levels positively correlated with SRF, cIAP2 and PCNA levels in cardiac fibroblasts from Spontaneously Hypertensive Rats. To conclude, DDR2-mediated ERK1/2MAPK activation facilitates coordinated regulation of cell survival and cell cycle progression in cardiac fibroblasts via SRF.

**New & Noteworthy:** Relative resistance to apoptosis and the ability to proliferate and produce a collagen-rich scar enable cardiac fibroblasts to play a central role in myocardial response to injury. This study reports novel findings that mitogen-stimulated cardiac fibroblasts exploit a common regulatory mechanism involving collagen receptor (DDR2)-dependent activation of ERK1/2 MAPK and SRF to achieve coordinated regulation of apoptosis resistance and cell cycle progression, which could facilitate their survival and function in the injured myocardium.

## Introduction

Cardiac fibroblasts, the principal stromal cells in the myocardium and a major source of matrix proteins, pro-inflammatory cytokines and pro-fibrotic factors, play an important role in wound healing following cardiac muscle damage (12). Unlike cardiac myocytes, cardiac fibroblasts retain their replicative capacity throughout adult life and are relatively resistant to pro-apoptotic signals such as Angiotensin II, TNF-α, hypoxia and oxidative stress that do not favor the survival of cardiac myocytes in the injured myocardium (22). In the altered cytokine milieu of the damaged heart, marked by progressive loss of functional cardiomyocytes, normally quiescent cardiac fibroblasts survive and undergo phenotypic transformation into active myofibroblasts that migrate to the site of injury, proliferate and produce matrix components to replace the lost tissue with a scar that maintains the structural and functional integrity of the heart (22). The relative resistance of cardiac fibroblasts to death signals that prevail in the myocardium *post injury* and their ability to proliferate and produce a collagen-rich scar are key determinants of their central role in acute wound healing.

Surprisingly, however, while there has been a great deal of interest in the regulation of collagen turnover in cardiac fibroblasts (10), only a limited number of studies have addressed the molecular mechanisms that determine cell survival and cell cycle progression in cardiac fibroblasts (13, 35, 38, 44, 46, 48, 49). Moreover, it is of obvious interest to ascertain whether the cell survival and cell cycle pathways are under distinct regulatory mechanisms or are co ordinately regulated in cardiac fibroblasts to achieve an efficient coupling of the two processes that are indispensable for optimal myocardial recovery from an acute insult.

We had demonstrated an obligate role for Discoidin Domain Receptor 2(DDR2), a mesenchymal cell-specific collagen receptor tyrosine kinase localized predominantly in fibroblasts in the heart, in collagen type I expression in cardiac fibroblasts and wound healing in response to Ang II, which points to the centrality of DDR2 in cardiac fibroblast response to injury (14). DDR2 has been implicated in a variety of fundamental cellular processes such as proliferation, survival and differentiation (30, 32, 60). While there are sporadic reports on the link between DDR2 and cell proliferation, demonstrated mostly in DDR2 null mice and in cancer cells (17, 28, 36, 42, 43), the direct involvement of DDR2 in the cell cycle machinery *per se* and the relevant signalling pathways activated by it remain, to the best of our knowledge, poorly defined. Further, the protective role of DDR2 in cells exposed to ambient stress and the relevant effectors and mechanisms involved remain largely obscure.

The present study provides robust evidence for the first time that mitogen-stimulated cardiac fibroblasts exploit a common regulatory mechanism involving collagen receptor (DDR2)-dependent activation of ERK1/2 MAPK and Serum Response Factor (SRF) to achieve coordinated regulation of apoptosis resistance and cell cycle progression, which would facilitate their survival and function in the injured myocardium.

## Materials and Methods

### Materials

Angiotensin II, CCG-1423, Candesartan cilexetil, CCG-1423, P D-98059 and M199 were obtained from Sigma-Aldrich (St. Louis, MO, USA). WRG-28 was obtained from MedChemExpress (Monmouth, NJ, USA). Random primers, reverse transcriptase, RNAase inhibitor, dNTPs and were obtained from Promega (Madison, WI, USA). The PureLink RNA isolation kit and Lipofectamine 2000 were from Invitrogen (Carlsbad, CA, USA). The Low cell# ChIPkit protein A × 48 was from Diagenode (Denville, NJ, USA). TaqMan probes for mRNA expression and Chemiluminescence western blot detection reagent were from Thermo Fisher Scientific (Waltham, MA, USA). DDR2 and control siRNAs were from Ambion (Foster City, CA, USA). SRF siRNAs were custom-designed from Eurogentec (Liege, Belgium). Signal Silence ® P44/42 MAPK (ERK1/2) siRNA was obtained from Cell Signaling Technology (Danvers, MA, USA). The rat DDR2/CD167b Gene ORF cDNA clone expression plasmid was obtained from Sino Biologicals (Beijing, China) and Native ORF cIAP-2 clone in pCMV vector was purchased from Origene, Rockville, USA. Opti-MEM and fetal Calf serum (FCS) were from GIBCO (Waltham, MA, USA). All cell culture ware was purchased from BD Falcon (Corning, NY USA). Primary antibodies against DDR2, p27KIP1, extracellular signal-regulated kinase 1/2 (ERK1/2) mitogen-activated protein kinase (MAPK), Total FoxO3a, Phospho-FoxO3a and cleaved-caspase-3 were obtained from Cell Signaling Technology (Danvers, MA, USA). The primary antibodies for cIAP2, Cyclin D1, Cyclin E, Skp2, procaspase-3 and Phospho-tyrosine were from Santa Cruz Biotechnology (Dallas, TX, USA). Rb and Phospho-Rb antibodies were purchased from Elabscience (Houston, TX, USA). Primary antibodyagainst SRF (Serum Response Factor) was obtained fromThermo Fisher Scientific (Waltham, MA, USA). Loading control β-Actin antibody was obtained from Sigma-Aldrich, (St. Louis, MO, USA). All antibodies were used after dilution (1:1000), except SRF (1:50) and FoxO3a (1:50) for chromatin immunoprecipitation (ChIP). XBT X-ray Film was from Carestream (Rochester, NY, USA). The study on rats was approved by the Institutional Animal Ethics Committees of SCTIMST (B form No: SCT/IAEC-233/AUGUST/2017/94 and SCT/IAEC-268/FEBRUARY/2018/95).

### Methods

#### Isolation of cardiac fibroblasts

Cardiac fibroblasts were isolated from young adult male Sprague–Dawley rats (2–3 months old) as described earlier (27). Sub-confluent cultures of cardiac fibroblasts from passage 2 or 3 were used for the experiments. Cells were serum-deprived for 24 h prior to treatment with 1μM Ang II, 25μM H2O2 or 10% Fetal Calf Serum (mitogen). Cells were pre-incubated with 10μM Candesartan (AT1 receptor antagonist) for 1 h before the addition of 25μM H2O2 in the appropriate group.

Cardiac fibroblasts were also isolated from 6 month-old male Wistar and Spontaneously Hypertensive Rats (SHR) as described earlier (27). Cells were collected after a brief wash 2.5h post initial plating to obtain cardiac fibroblasts. These cells were collected and processed further to analyse the expression of various genes and proteins.

#### Quantitative reverse transcription-polymerase chain reaction (RT-qPCR) analysis

Sub-confluent cultures of cardiac fibroblasts were subjected to the indicated treatments and total RNA was isolated using the PureLink RNA isolation kit (Invitrogen), according to the manufacturer’s instructions. Following DNase I treatment, 2μg of total RNA was reverse transcribed to cDNA with random primers and M-MLV reverse transcriptase. TaqMan RT qPCR analysis was carried out using the ABI prism 7500 Sequence Detection System (Applied Biosystems, CA, USA) with specific FAM-labeled probes for cIAP2(Birc2) (Assay ID: Rn00572734_m1), and VIC-labeled probes for β-actin (Rn00667869_m1). PCR reactions were performed under the following thermal cycling conditions: 95°C for 10 min followed by 40 cycles of denaturation at 95°C for 15 s and annealing/extension at 60°C for 1 min. Gene expression was quantified using *C*_T_ values. mRNA expression was normalized to that of β-actin. The relative fold-change in target mRNA levels of treated versus control was quantified using the 2^-ΔΔ*C*t^ method.

#### Western blot analysis

Sub-confluent cultures of cardiac fibroblasts in serum-free M199 were treated with Ang II (1μM), or 10% Fetal Calf Serum (mitogen) and relative protein abundance was determined by western blot analysis following standard protocols, with β-actin as loading control. Enhanced chemiluminescence reagent was used to detect the proteins with X-ray Film.

#### RNA interference and over-expression

Cardiac fibroblasts at passage 3 were seeded on 60mm dishes at equal density. After 24h, the cells were incubated in Opti-MEM for 5–6h with Ambion pre-designed Silencer-Select siRNA, custom-designed siRNA from Eurogentech or scrambled siRNA (control siRNA) at the given concentrations (10nM for DDR2, 20nM for SRF and ERK1/2 MAPK) and Lipofectamine 2000 (8μl).

Constitutive expression of DDR2 and cIAP2 was achieved under the control of a CMV promoter. The DDR2 and cIAP2 plasmids were verified by restriction mapping. For over expression, the plasmid vector for DDR2 or cIAP2 (1μg/μl) was transfected using Lipofectamine 2000. The plasmid alone without the cDNA insert was used as control for transfection. Following a post-transfection recovery phase in M199 with 10% FCS for 12h, the cells were serum-deprived for 24h and then treated with Ang II (1μM) or 10% FCS or H_2_O_2_ (25μM) for the indicated durations. Cell lysates were prepared in Laemmli sample buffer, denatured and used for western blot analysis.

#### Chromatin Immunoprecipitation (ChIP) assay

The ChIP assay was performed with the Low Cell Number ChIP kit, according to the manufacturer’s protocol. Briefly, after treatment of cardiac fibroblasts with 1μM Ang II or 10% FCS for 30 min, the cells were cross-linked with 1% formaldehyde, lysed and sonicated in a Diagenode Bioruptor to generate ~600 bp DNA fragments. The lysates were incubated with anti-SRF or anti-FoxO3a antibody overnight at 4°C with rotation. Immune complexes were precipitated with protein A-coated magnetic beads. After digestion with proteinase K to remove the DNA-protein cross-links from the immune complexes, the DNA was isolated and subjected to PCR using primers for the specific promoter regions. In samples immunoprecipitated with the SRF antibody, the cIAP2 proximity region was amplified using FP-5’-AAGGGGTAAAAGATTTGAGG-3’ and RP-5’-CTATCAACATTGGAGACCAAG-3’, which amplifies a region containing CAARG element (CCTTAAAAGG) 1982bp downstream of the start site in the first intron. Skp2 proximity region was amplified using FP-5’ AGGACAGCCAAGACTACAAAG 3’ and RP-5’ ATAACAGGCAAATGACCCTTC 3’, which amplifies a region containing CAARG element (CCAAAAAAGG) 10,819 bp downstream of start site in fifth intron. In samples immunoprecipitated with the FoxO3a antibody, p27 proximity region was amplified with FP-5’ GAGACGTGGGGCGTAGAATAC 3’ and RP-5’ ATTCGGGGAACCGTCTGAAAC 3’, which amplifies a region containing FOXO consensus element (CAAAACAA) 1,016bp downstream of the start site in the second exon. DNA isolated from an aliquot of the total sheared chromatin was used as loading control for PCR (input control). ChIP with a non specific antibody (normal rabbit IgG) served as negative control. The PCR products were subjected to electrophoresis on a 2% agarose gel.

#### Effect of serum from SHR on normal cardiac fibroblasts

Preparation of serum: Blood was collected from the descending aorta of anesthetized rats, and serum was separated by centrifugation. For experiments with cardiac fibroblasts, serum was filtered through 0.22μm membrane and used fresh. Cardiac fibroblasts from Wistar rats were exposed to serum from Wistar rats and SHR to achieve a final serum concentration of 10% in Medium M199.

#### Flow cytometry

##### Annexin V-PI staining

Cells were trypsinized, washed twice with cold PBS and re-suspended in 1X Binding Buffer at a concentration of 1 × 10^6^ cells/ml. 5 μl of FITC Annexin V and 5 μl PI were added to 1 × 10^5^ cells/100ul in a 5 ml culture tube, gently vortexed and incubated for 15 min at RT (25°C) in the dark. Cells were analyzed by flow cytometry after adding 400 μl of 1X Binding Buffer to each tube. Unstained cells, cells stained with only FITC Annexin V and cells stained with only PI were used as controls to set up compensation and quadrants. Flow cytometry was perfomed using BD FACSJazz^™^ Cell Sorter.

##### Cell cycle analysis

Cells were trypsinized, washed twice with cold PBS, re-suspended in 500ul PBS, fixed with an equal volume of 70% ethanol and stored at 4°C overnight. The cells were pelleted, washed twice in 1xPBS, re-suspended in 500μl 1xPBS and incubated at room temperature for 30 minutes with 50ug of RNase A. Following incubation with 200ul of Propidium Iodide in the dark for 15 min, the cells were analyzed by flow cytometry on BD FACSJazz^™^ Cell Sorter.

##### Nuclear and Cytoplasmic isolation

Cells were harvested by trypsinization following specific treatments, washed in ice-cold PBS thrice and the final cell pellet was divided into two fractions, one for western blot analysis and the other for cell fractionation. Cell fractionation was performed using NE-PER Nuclear and Cytoplasmic Extraction Reagent (Thermo Scientific), following the kit protocol. The cytoplasmic and nuclear extracts obtained were subjected to western blot analysis using HDAC1 as nuclear marker and GAPDH as cytoplasmic marker.

##### Immunocytochemistry

Cells were cultured on chamber slides and subjected to various treatments. After fixation, permeabilization and blocking, the cells were incubated with the FoxO3a (total) primary antibody (1:200) for 1h at RT, and with respective Alexa Fluor 488-conjugated goat anti rabbit secondary antibody (1:200) for 1h. Actin was stained with phalloidin conjugated with rhodamine along with secondary antibody and incubated at room temperature. Nuclei were counterstained with Hoechst 33258 for 15Ↄmin and imaged using fluorescence microscope (ZEISS AXIO Imager.A2).

##### Immunoprecipitation of DDR2

Cells were harvested following specified treatments at different time-points and lysed in TNN IP buffer (50 mmol/L Tris-HCl (pH 7.5), 150 mmol/L NaCl, 0.5% NP40, protease and phosphatase inhibitors without EDTA). The protein was estimated and 100ug protein was immunoprecipitated with 2ug DDR2 antibody by incubation at 40C overnight on spin rotor at 30 RPM. An input sample was taken in one sample before adding antibody to evaluate the efficiency of pull-down. The DDR2 antibody was pulled down using Protein A magnetic beads. The pull-down efficiency was analysed by western blotting, loading the input sample, unbound fraction and pull-down fraction (Figure 10A). Further, the lysates were analysed by western blotting for presence of Phospho-tyrosine and Total DDR2 levels in each sample.

#### Statistical analysis

Data are expressed as Mean±SE. GraphPad Prism 6 Software was used for the graphs and statistical tests. Student’s *t* test (unpaired, 2-tail) was done for comparisons involving 2 groups and two-way ANOVA was done for comparisons involving more than 2 groups with two variables. Tukey’s multiple comparisons test was also done. p<0.05 was considered significant. The in vitro data presented are representative of 3 or 4 independent experiments (n=3 or 4). Three age-matched (6 months) male SHR and Wistar rats were used in the in vivo experiments (n=3).

## Results

### DDR2 mediates Ang II-stimulated expression of anti-apoptotic cIAP2 in cardiac fibroblasts

The intra-cardiac generation of Ang II is enhanced following myocardial injury, which exposes the different cell types in the heart to its pleiotropic actions (55). Paradoxically, while Ang II is reported to induce apoptosis in cardiac myocytes (2, 24), it promotes cardiac fibroblast activation *post injury* and is a potent pro-fibrotic factor with marked stimulatory effect on collagen expression in cardiac fibroblasts (55). These observations prompted us to explore the possible pro-survival role of Ang II in cardiac fibroblasts and delineate the underlying mechanisms, focusing specifically on the role of DDR2.

First, we examined the effect of Ang II on the expression of cellular Inhibitor of Apoptosis, cIAP2, which was earlier shown by us to play a key role in cardiac fibroblast resistance to oxidative stress (48). Ang II induced a 3-fold increase in cIAP2 mRNA at 6h (Figure 1A) and 2-fold increase in cIAP2 protein expression at 12h post-treatment (Figure 1B), determined by RT-qPCR and western blotting, respectively.

**Figure 1:**
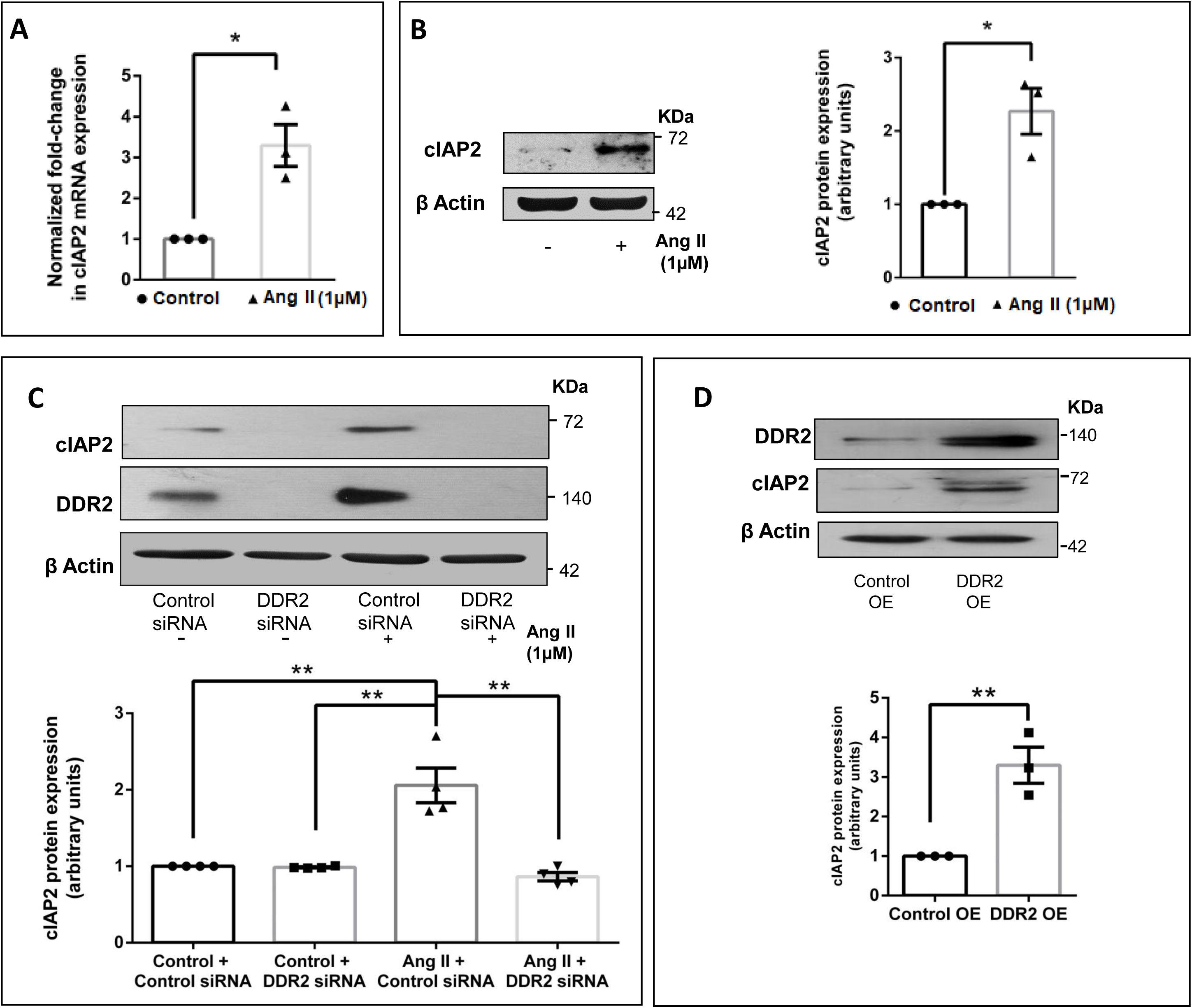
DDR2 mediates Ang II-stimulated expression of anti-apoptotic cIAP2 in cardiac fibroblasts. Sub-confluent quiescent cultures of cardiac fibroblasts were stimulated with Ang II (1μM). **(A)** cIAP2 mRNA levels were determined by Taqman Real-time PCR analysis at 6h of Ang II treatment. β-actin served as the endogenous control. Significance was determined by Student’s t test, *p< 0.05 vs. control. (**B**) Protein was isolated at 12h of Ang II treatment and subjected to western blot analysis for detection of cIAP2, with β-actin as loading control. Significance was determined by Student’s t test, *p< 0.05 vs. control. (**C**) RNAi-mediated silencing of DDR2 confirmed its role in regulating cIAP2 gene expression in Ang II-stimulated cardiac fibroblasts. Cardiac fibroblasts were transiently transfected with DDR2 siRNA (5 pmol) or control (scrambled) siRNA prior to treatment with Ang II for 12h. cIAP2 protein expression was examined, with β-actin as loading control. Validation of DDR2 silencing is also shown. Significance was determined by two-way ANOVA (**Tukey’s** multiple comparisons test, **p< 0.01, comparisons as depicted in the Figure) (**D**) Cardiac fibroblasts were transfected with DDR2 cDNA over-expression plasmid (DDR2OE) (with empty vector control, Control OE), post-revival, the cells were serum-deprived for 24h. Cells were collected and cIAP2 protein expression was examined, with β-actin as loading control. Significance was determined by Student’s t test, **p< 0.01 vs Control OE. Validation of DDR2 over-expression is also shown. Data are representative of 3 or 4 independent experiments, n=3 or 4. Mean ± SEM (Standard Error of Mean).

Notably, DDR2 knockdown in cardiac fibroblasts using specific siRNA prevented Ang II stimulated cIAP2 expression (Figure 1C), showing that DDR2 mediates the stimulatory effect of Ang II on cIAP2 expression. Further, DDR2 over-expression in un-stimulated cells enhanced cIAP2 expression (Figure 1D), clearly demonstrating its role in cIAP2 regulation.

### DDR2-dependent ERK1/2 MAPK activation acts via Serum Response Factor to transcriptionally up-regulate cIAP2 expression in Ang II-stimulated cardiac fibroblasts

Since bioinformatics analysis of the cIAP2 promoter region revealed binding sites for Serum Response Factor (SRF), we probed its possible involvement in the regulation of Ang II dependent cIAP2 expression. We found that inhibition of Myocardin-Related Transcription Factor A/B (MRTF-A/B), a cofactor of SRF (45, 62), using CCG-1423 that inhibits both the MRTF isoforms (20) led to downregulation of cIAP2 protein expression (Figure 2A) without affecting total SRF and DDR2 levels (Figure 2A), indicating MRTF involvement in the cellular response to Ang II. Further, SRF knockdown with specific siRNA was found to abolish cIAP2 in Ang II-stimulated cells, confirming its role in cIAP2 expression (Figure 2B). Importantly, while SRF knockdown did not affect DDR2 expression (Figure 2B), knockdown of DDR2 down-regulated SRF levels (Figure 2C), confirming that DDR2 regulates cIAP2 via SRF in Ang II-stimulated cells. Further, chromatin immunoprecipitation assay (ChIP) demonstrated SRF binding to the promoter region of cIAP2 in Ang II-treated cells, which was attenuated in DDR2-silenced cells (Figure 2D).

**Figure 2:**
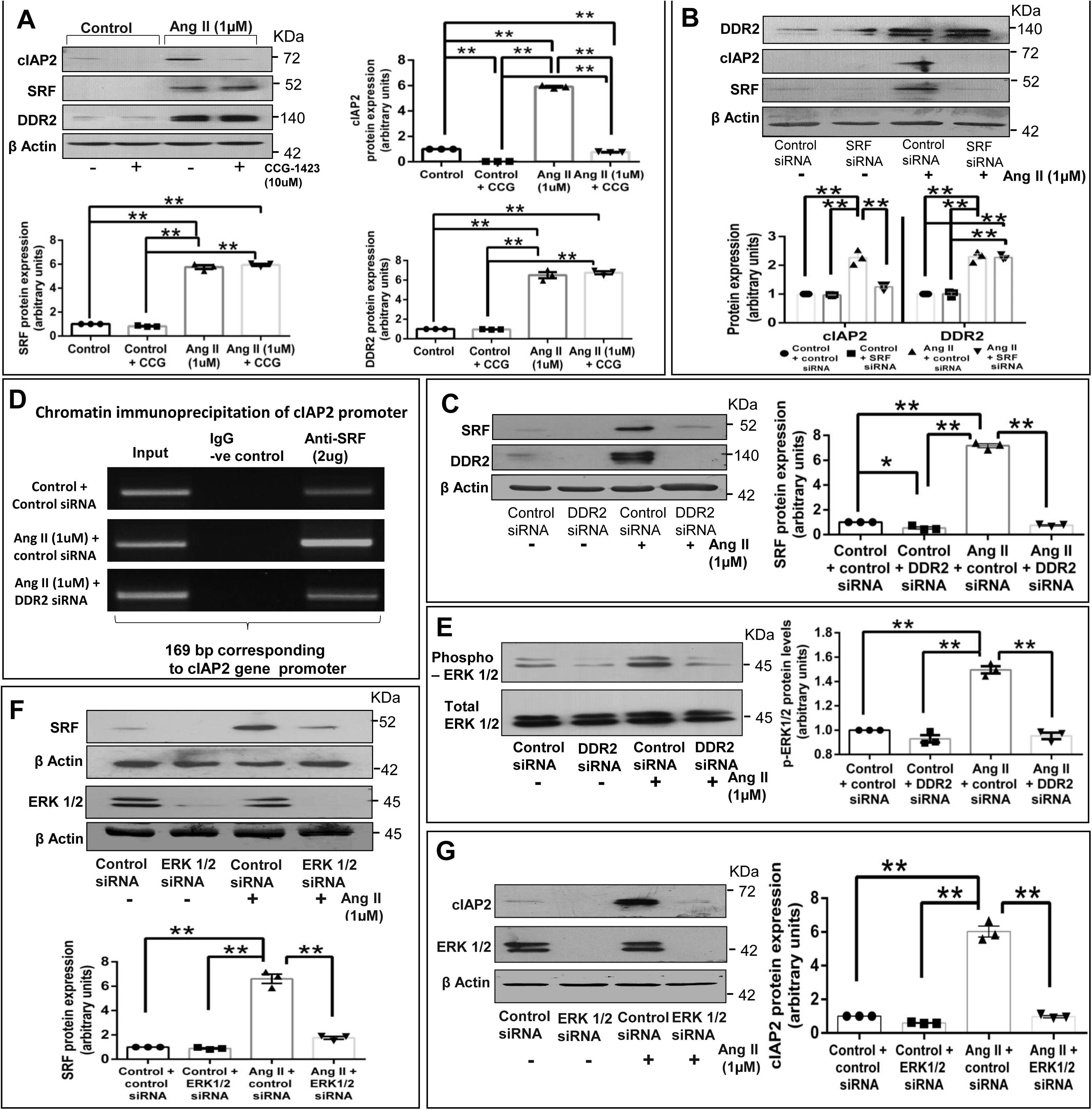
DDR2-dependent ERK1/2 MAPK activation acts via SRF to transcriptionally upregulate cIAP2 expression in Ang II-stimulated cardiac fibroblasts. (**A**) Sub-confluent quiescent cultures of cardiac fibroblasts were pre-treated with CCG-1423(10μM) for 45 min prior to treatment with Ang II (1μM) for 12h. cIAP2, SRF and DDR2 protein expression was examined by western blot analysis, with β-actin as loading control. Significance was determined by two-way ANOVA (Tukey’s multiple comparisons test, **p< 0.01, comparisons as depicted in the Figure). (**B**) Sub-confluent quiescent cultures of cardiac fibroblasts were transiently transfected with SRF siRNA (10 pmol) or control (scrambled) siRNA prior to treatment with Ang II (1μM) for 12h. cIAP2 and DDR2protein expression was examined by western blot analysis, with β-actin as loading control. Significance was determined by two-way ANOVA (Tukey’s multiple comparisons test, **p< 0.01, comparisons as depicted in the Figure). (**C-E**) Sub-confluent quiescent cultures of cardiac fibroblasts in M199 were transiently transfected with DDR2 siRNA (5pmol) or control (scrambled) siRNA prior to treatment with Ang II (1μM). (**C**) Cells were collected at 12h post-Ang II treatment and SRF protein expression was examined by western blot analysis, with β-actin as loading control. Significance was determined by two-way ANOVA (Tukey’s multiple comparisons test, *p< 0.05 and **p< 0.01, comparisons as depicted in the Figure). (**D**) Cells were collected at 30 mins post-Ang II treatment and chromatin was immunoprecipitated by anti-SRF antibody followed by PCR amplification and analysed on a 2% agarose gel for presence of 169bp region of cIAP2 gene promoter (region amplified is specified in methods section) (**E**) Cells were collected at 12h post-Ang II treatment and Phospho-ERK1/2 protein level was examined by western blot analysis, with Total ERK1/2 level as loading control. Significance was determined by two-way ANOVA (Tukey’s multiple comparisons test, **p< 0.01, comparisons as depicted in the Figure). (**F-G**) Subconfluent quiescent cultures of cardiac fibroblasts in M199 were transiently transfected with ERK1/2 siRNA (10pmol) or control (scrambled) siRNA prior to treatment with Ang II (1μM). (**F**) Cells were collected at 12h post-Ang II treatment and SRF protein expression was examined by western blot analysis, with β-actin as loading control. Significance was determined by two-way ANOVA (Tukey’s multiple comparisons test, **p< 0.01, comparisons as depicted in the Figure). ERK1/2 knockdown validation is also shown. (**G**) Cells were collected at 12h post-Ang II treatment and cIAP2 protein expression was examined by western blot analysis, with β-actin as loading control. Significance was determined by two-way ANOVA (Tukey’s multiple comparisons test, **p< 0.01, comparisons as depicted in the Figure). ERK1/2 knockdown validation is shown. Data are representative of 3 independent experiments, n=3. Mean ± SEM (Standard Error of Mean).

As previously reported by us (14), we observed DDR2-dependent activation of ERK1/2 MAPK in Ang II-stimulated cells (Figure 2E). Further, ERK1/2 MAPK knockdown down regulated Ang II-stimulated SRF expression and cIAP2 expression (Figure 2F and Figure 2G), showing that DDR2 increases cIAP2 levels via ERK1/2 MAPK-dependent SRF in Ang II-stimulated cells.

### DDR2-dependent cIAP2 expression protects cardiac fibroblasts against oxidative damage

We had reported earlier that cIAP2, induced in response to H_2_O_2_, protects cardiac fibroblasts against oxidative damage (48). Additionally, we had also shown that H_2_O_2_ treatment promotes Ang II production in cardiac fibroblasts (1). Pursuing these observations, we found that blocking the Ang II receptor, AT1, with candesartan attenuated cIAP2 expression (Figure 3A.) and promoted apoptosis in H_2_O_2_-treated cells (Figures 3B and Supplementary Figure S1A https://doi.org/10.6084/m9.figshare.11371227.v1), showing that Ang II produced in response to H_2_O_2_ is responsible for cIAP2 induction and protection against oxidative damage. Interestingly, H_2_O_2_ treatment also enhanced DDR2 and SRF expression, which was abolished by candesartan (Figure 3A). Further, we found that DDR2 and SRF knockdown in H_2_O_2_-treated cardiac fibroblasts reduced cIAP2 expression (Figures 3C and 3D). Knockdown of DDR2 and SRF was found to increase levels of cleaved-caspase-3 (Figures 3E and 3F) and promote cell death in H_2_O_2_-treated cardiac fibroblasts (Figures 3G and Supplementary Figure S1B https://doi.org/10.6084/m9.figshare.11371227.v1). Interestingly, over-expression of cIAP2 gene was found to abolish the activation of apoptosis in DDR2-silenced cardiac fibroblasts exposed to H_2_O_2_ (Figure 3H). Together, these data point to the pro-survival role of Ang II in cardiac fibroblasts under oxidative stress by a mechanism involving DDR2-dependent transcriptional up-regulation of cIAP2 expression by SRF.

**Figure 3:**
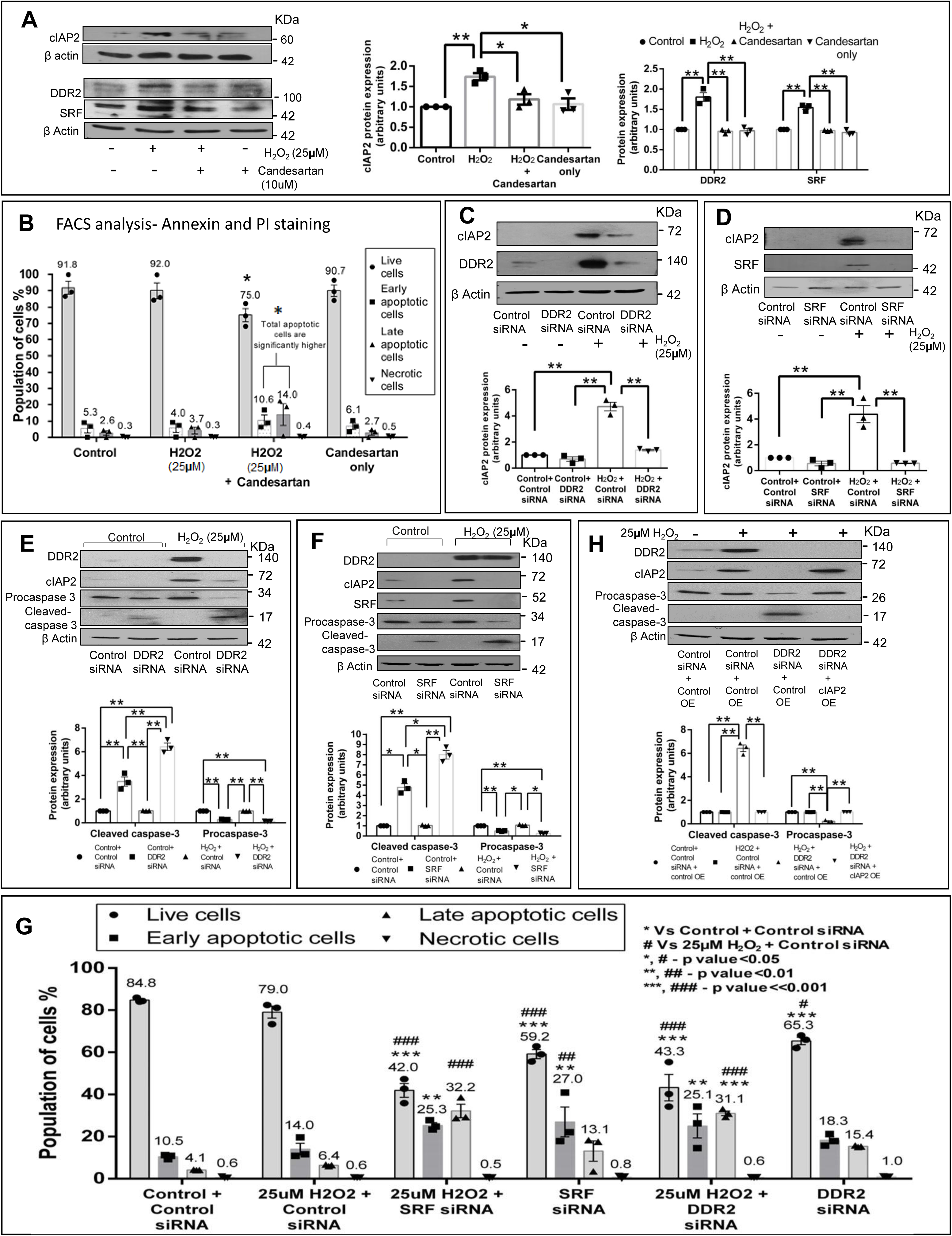
DDR2-dependent cIAP2 expression protects cardiac fibroblasts against oxidative damage. (**A-B**) Effect of AT1 receptor antagonist, Candesartan, on H_2_O_2_-treated cardiac fibroblasts was analysed. Four sets of sub-confluent cultures of cardiac fibroblasts were serum-deprived for 24h: i) no treatment (control), ii) 25μM H_2_O_2_ iii) pre-incubated with 10μM candesartan (AT1 receptor antagonist) for 1 h and was treated with 25μM H_2_O_2_ iv) 10μM candesartan alone. (**A**) Cells were collected 12h post-H_2_O_2_ addition and analysed by western blot for the expression of cIAP2, DDR2 and SRF, with β-actin as loading control. Significance was determined by two-way ANOVA (Tukey’s multiple comparisons test, *p< 0.05 and **p< 0.01, comparisons as depicted in the Figure). (**B**) Cells were collected 8h post-H_2_O_2_ addition and analysed by flow cytometry for annexin/PI uptake and represented as percentage of cells. Significance was determined by two-way ANOVA (Tukey’s multiple comparisons test, *p< 0.05, Live cells in 25μM H_2_O_2_ + 10μM candesartan vs control or 25μM H_2_O_2_ or candesartan alone). *p<0.05 Apoptotic cells (early apoptotic + late apoptotic cells) in 25μM H_2_O_2_ + 10μM candesartan vs control or 25μM H_2_O_2_ or candesartan alone. Necrotic cells were insignificant in number (see also Supplementary Figure S1 A https://doi.org/10.6084/m9.figshare.11371227.v1). (**C-G**) Effect of DDR2 or SRF gene silencing on H_2_O_2_-treated cardiac fibroblasts was analysed. Sub-confluent quiescent cultures of cardiac fibroblasts in M199 were transiently transfected with DDR2 siRNA (5pmol) or SRF siRNA(10pmol) with respective control (scrambled) siRNA prior to treatment with 25μM H_2_O_2_. (**C and D**) Cells were collected 12h post-H_2_O_2_ addition and analysed by western blot for the expression of cIAP2, with β-actin as loading control. Significance was determined by two-way ANOVA (Tukey’s multiple comparisons test, **p< 0.01, comparisons as depicted in the Figure). (**E and F)** Cells were collected 8 h post-H_2_O_2_ addition and analysed by western blot for the expression levels of cleaved-caspase 3 and procaspase 3, with β-actin as loading control. Validation of DDR2 and SRF knockdown is also shown along with corresponding cIAP2 levels in the same blot. Significance was determined by two-way ANOVA (Tukey’s multiple comparisons test, *p<0.05 and **p< 0.01 (comparisons as depicted in the Figure). (**G**) Cells were collected 8h post-H_2_O_2_ addition and analysed by flow cytometry for annexin/PI uptake and represented as percentage of cells. Significance was determined by two-way ANOVA (Tukey’s multiple comparisons test, * vs control + control siRNA, # vs 25μM H_2_O_2_+ control siRNA) *, # - p < 0.05; **, ## - p< 0.01 and ***, ### -p<<0.001). See also Supplementary Figure S1 B https://doi.org/10.6084/m9.figshare.11371227.v1. (**H**) Effect of cIAP2 overexpression in DDR2-silenced and H_2_O_2_-treated cardiac fibroblasts was analysed. Sub-confluent quiescent cultures of cardiac fibroblasts in M199 were transiently transfected with DDR2 siRNA (5pmol) alone and with cIAP2 cDNA over-expression plasmid (cIAP2 OE) with respective scrambled (control) siRNA and plasmid alone (Control OE) prior to treatment with 25μM H_2_O_2_. Cells were collected 8h post-H_2_O_2_ addition and analysed by western blot for the expression levels of cleaved-caspase 3 and procaspase 3, with β-actin as loading control. Validation of DDR2 knockdown and cIAP2 overexpression are also shown in the same blot. Significance was determined by two-way ANOVA (Tukey’s multiple comparisons test, **p< 0.01 (comparisons as depicted in the Figure). Data are representative of 3 independent experiments, n=3. Mean ± SEM (Standard Error of Mean).

#### An obligate role for DDR2 in G1-S transition in cardiac fibroblasts via transcriptional and post-translational regulation of p27

To evaluate the role of DDR2 in G1-S transition, we employed a combination of gene knockdown and over-expression approaches and examined cell cycle status in relation to DDR2 expression. 10% Fetal Calf Serum was used to obtain a robust mitogenic effect, which would facilitate an unequivocal assessment of the regulatory role of DDR2 in the cardiac fibroblast cell cycle. Our preliminary experiments had revealed that Ang II fails to induce significant changes in PCNA and cyclin D1 expression (Figure 4A), consistent with earlier reports on the inability of Ang II to directly trigger mitogenesis in cardiac fibroblasts through activation of the cyclin-dependent pathway (3). In fact, H_2_O_2_, which generates Ang II in cardiac fibroblasts, also failed to induce significant changes in PCNA and cyclin D1 expression (Figure 4A).

**Figure 4:**
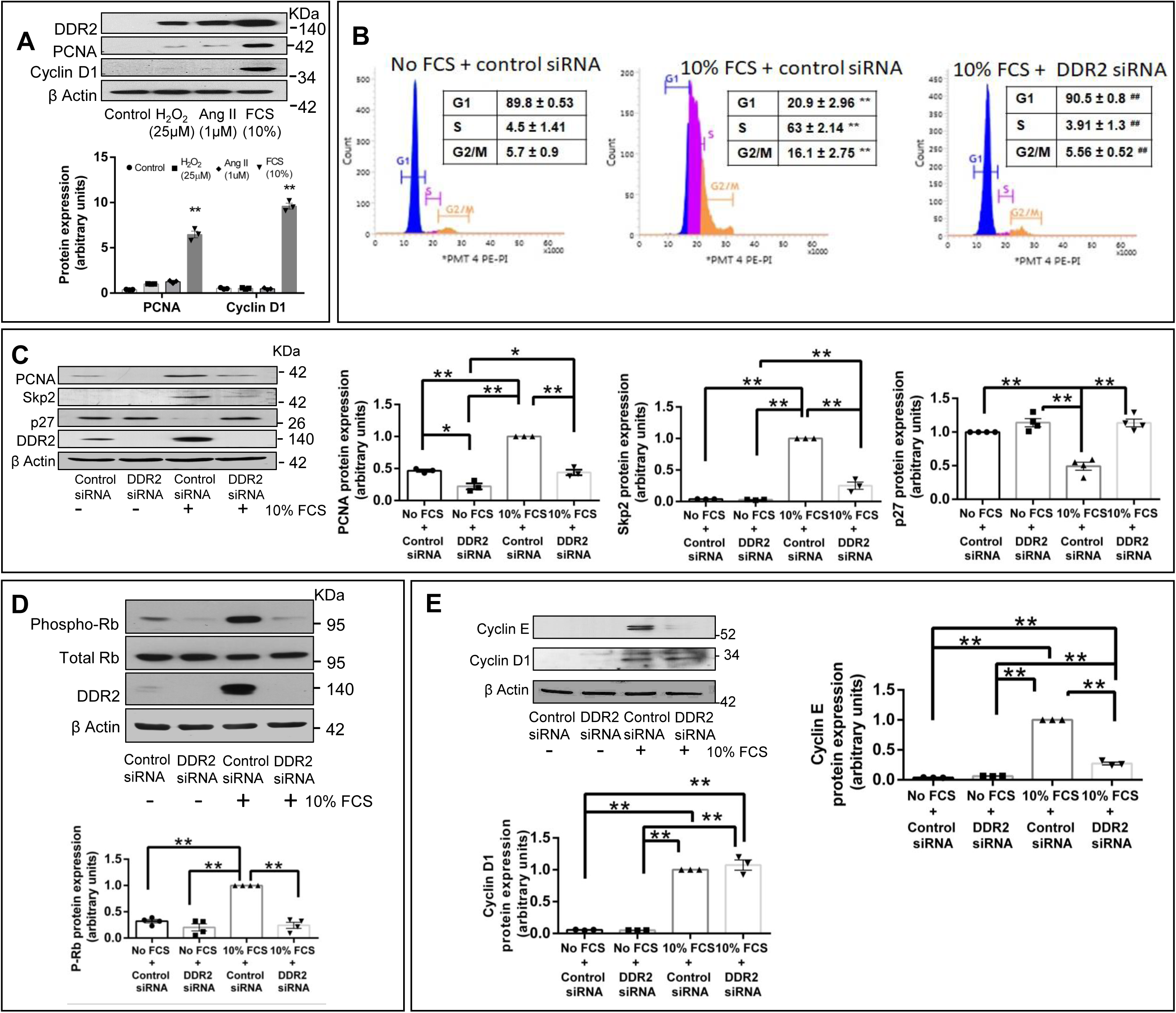
DDR2 knockdown in mitogen-stimulated cardiac fibroblasts inhibits G1-S transition. (**A)** Sub-confluent cultures of cardiac fibroblasts were serum deprived for 24h prior to treatment with Control (fresh M199 medium treatment), 25μM H_2_O_2_, Ang II (1μM) and 10% Fetal Calf Serum. Post treatment, cells were collected at 8h and analysed by western blotting for expression levels of PCNA and cyclin D1, with β-actin as loading control. Significance was determined by Student’s t test, (each treatment vs Control) **p<0.01 10% FCS vs Control. 25μM H_2_O_2_ and Ang II (1μM) are not significant *(**B-E**)* Sub-confluent cultures of cardiac fibroblasts were transfected with DDR2 siRNA (or control siRNA) and, following revival for 12h in 10% serum-supplemented medium, the cells were serumdeprived for synchronization. Post-synchronization, cells were exposed to 10% Fetal Calf Serum (10% FCS). (**B**) Cells were collected at 14h for flow cytometric analysis of G1-S transition, showing distribution of cells in each phase in percentage. Significance was determined by one-way ANOVA (Tukey’s multiple comparisons test, **p<0.01 vs no FCS + control siRNA, ## p< 0.01 vs 10% FCS + control siRNA). (**C**) Cells were collected at 8h and analysed by western blotting for expression levels of PCNA, Skp2 and p27, with β-actin as loading control. Significance was determined by two-way ANOVA (Tukey’s multiple comparisons test, *p<0.05 and **p< 0.01, comparisons as depicted in the Figure). (**D**) Cells were collected at 8h and analysed by western blotting for expression levels of Phospho-Rb, with Total Rb as loading control. Significance was determined by two-way ANOVA (Tukey’s multiple comparisons test, **p< 0.01, comparisons as depicted in the Figure). (**E**) Cells were collected at 8h and analysed by western blotting for expression levels of Cyclin E and Cyclin D1, with β-actin as loading control. Significance was determined by two-way ANOVA (Tukey’s multiple comparisons test, ** p< 0.01, comparisons as depicted in the Figure). Data are representative of 3 or 4 independent experiments, n=3 or 4. Mean ± SEM (Standard Error of Mean).

#### DDR2 knockdown in mitogen-stimulated cardiac fibroblasts inhibits G1-S transition

Sub-confluent cultures of cardiac fibroblasts were transfected with DDR2 siRNA (or control siRNA) and, following revival for 12h in 10% serum-supplemented medium, the cells were serum-deprived for synchronization. Post synchronization, cells were exposed to mitogenic stimulation (10% Fetal Calf Serum) and collected at either 8h for analysis of various cell cycle regulatory molecules by western blotting or at 14h for flow cytometric analysis of G1-S transition. Flow cytometric analysis showed that DDR2 knockdown in mitogen-stimulated cells results in cell cycle arrest at the G1 phase (Figure 4B). Further, western blot analysis showed that DDR2 knockdown in mitogen-stimulated cells results in a significant reduction in the levels of Proliferating Cell Nuclear Antigen (PCNA), an S-phase marker (Figure 4C). Skp2 is an E3 ubiquitin ligase that targets various inhibitors of G1-S transition, including p27 that belongs to the CIP/KIP family of cyclin-dependent kinase (CDK) inhibitors, to facilitate cell cycle progression (5). We found that DDR2 knockdown attenuates Skp2 levels (Figure 4C), resulting in p27 induction (Figure 4C) and Rb hypophosphorylation (Figure 4D), culminating in G1 arrest of mitogen-stimulated cells. Further, cyclin E, an S-Phase cyclin, but not Cyclin D1, a G1-phase cyclin, was significantly reduced in DDR2-silenced, mitogen stimulated cells (Figure 4E). Together, these data suggest that DDR2 has an obligate role in G1-S transition via positive regulation of Skp2 and negative regulation of p27.

### Regulation of Skp2 by DDR2-dependent SRF

Based on bioinformatics analysis, the possible involvement of SRF in DDR2-dependent Skp2 expression was probed next. SRF knockdown was found to down-regulate Skp2 (Figure 5A), showing that SRF is involved in the regulation of Skp2. DDR2 and ERK1/2 MAPK knockdown in serum-stimulated cells led to down-regulation of SRF (Figure 5B, 5C). Since DDR2 is involved in ERK1/2 activation in serum-stimulated cells as well (Figure 5D), these data point to DDR2-dependent regulation of SRF via ERK1/2 MAPK. Together, the findings demonstrate a role for DDR2-dependent SRF activation in the regulation of Skp2. DDR2-dependent binding of SRF to the Skp2 gene promoter was confirmed by ChIP assay (Figure 5E).

**Figure 5:**
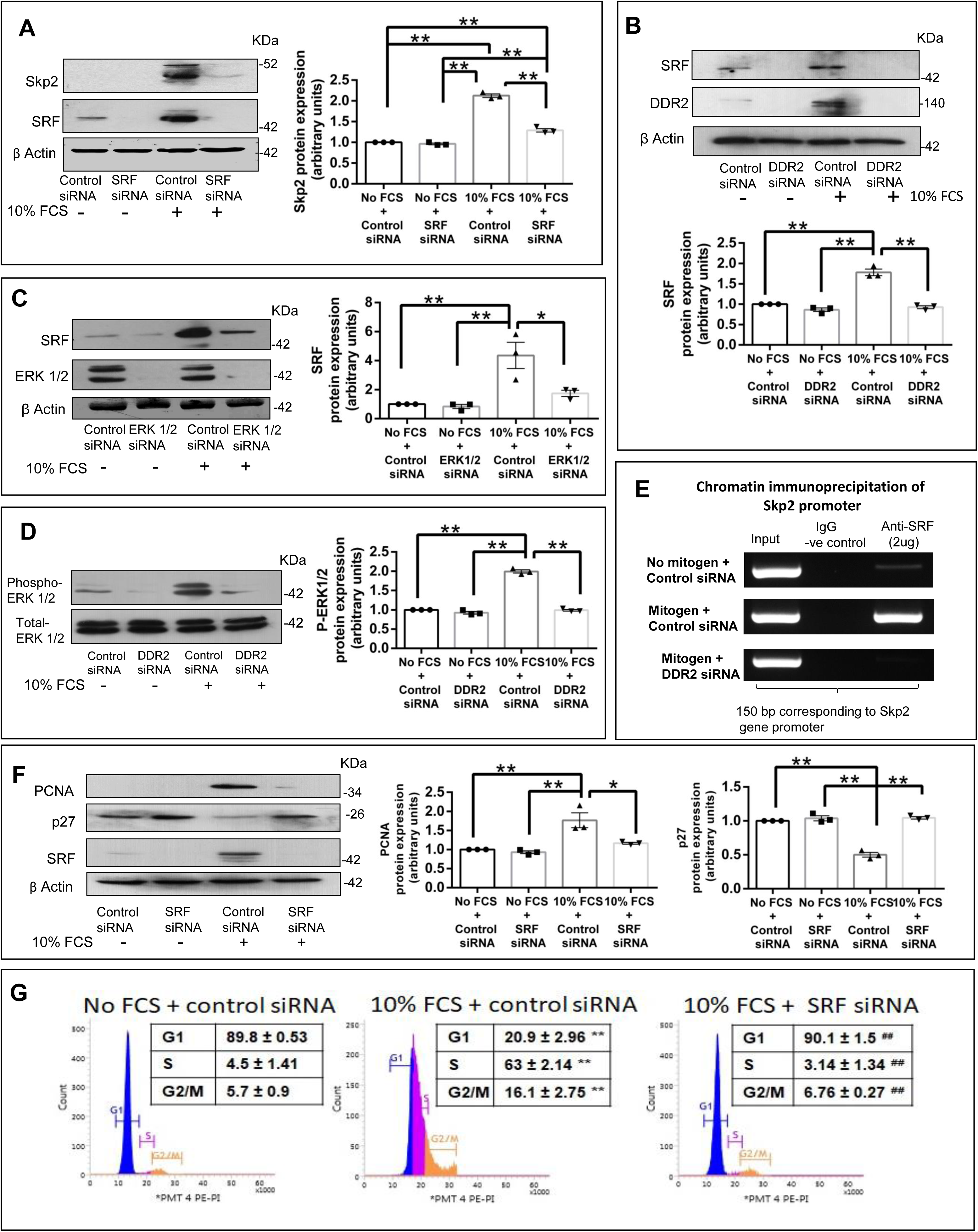
Regulation of p27: Post-translational regulation via SRF-dependent Skp2. (**A-G**) Sub-confluent cultures of cardiac fibroblasts were transfected with SRF siRNA (**A, F and G**), ERK1/2 siRNA (**C**) or DDR2 siRNA (**B, D** and **E**) (with Control siRNA) and, following revival for 12h in 10% serum-supplemented medium, the cells were serum-deprived for synchronization. Post-synchronization, the cells were exposed to 10% Fetal Calf Serum. (**A**) SRF siRNA transfected cells were collected at 8h and analysed by western blotting for expression levels of Skp2, with β-actin as loading control. Significance was determined by two-way ANOVA (Tukey’s multiple comparisons test, **p< 0.01 comparisons as depicted in the Figure). Validation of SRF knockdown is also shown. (**B**) DDR2 siRNA-transfected cells were collected at 8h and analysed by western blotting for expression levels of SRF, with β-actin as loading control. Significance was determined by two-way ANOVA (Tukey’s multiple comparisons test, **p<0.01, comparisons as depicted in the Figure). Validation of DDR2 knockdown is also shown. (**C**) ERK1/2 MAPK siRNA-transfected cells were collected at 8h and analysed by western blotting for expression levels of SRF, with β-actin as loading control. Significance was determined by two-way ANOVA (Tukey’s multiple comparisons test, *p<0.05 and **p<0.01, comparisons as depicted in the Figure). Validation of ERK1/2 MAPK knockdown is also shown. (**D**) DDR2 siRNA-transfected cells were collected at 8h and Phospho-ERK1/2 protein level was examined by western blot analysis, with Total ERK1/2 level as loading control. Significance was determined by two-way ANOVA (Tukey’s multiple comparisons test, **p< 0.01, comparisons as depicted in the Figure). (**E**) Sub-confluent cultures of cardiac fibroblasts were transfected with DDR2 siRNA (or control siRNA) and, following revival for 12h, cells were synchronized and exposed to 10% Fetal 30 Calf Serum. Cells were collected at 30 min and chromatin was immunoprecipitated using anti-SRF antibody; the image shows PCR amplified 150bp region of the Skp2 gene promoter on 2% agarose gel (region amplified is specified in methods section). (**F**) SRF siRNAtransfected cells were collected at 8h and analysed by western blotting for expression levels of PCNA and p27, with β-actin as loading control. Significance was determined by two-way ANOVA (Tukey’s multiple comparisons test, *p<0.05, **p< 0.01, comparisons as depicted in the Figure). (**G**) Cells were collected at 14h for flow cytometric analysis of G1-S transition, showing distribution of cells in each phase in percentage. Significance was determined by one-way ANOVA (Tukey’s multiple comparisons test, **p<0.01 vs no FCS + control siRNA, ## p< 0.01 vs 10% FCS + control siRNA). Data are representative of 3 independent experiments, n=3. Mean ± SEM (Standard Error of Mean).

### Regulation of p27 by DDR2

#### i) Post-translational regulation via SRF-dependent Skp2

Since Skp2 is known to post-translationally degrade p27 (5) and DDR2-dependent SRF activation regulates Skp2 (Figures 5A-E), we examined whether SRF silencing would affect p27 levels. SRF knockdown was found to induce p27 and reduce PCNA expression in mitogen-stimulated cells (Figure 5F) and cause cell cycle arrest at the G1 phase (Figure 5G). Together, these data suggest that SRF may transcriptionally regulate Skp2 expression and facilitate degradation of p27 and G1-S transition.

#### ii) Transcriptional regulation through modulating FoxO3a activity

Since p27 was found to be up-regulated in DDR2-silenced, mitogen-stimulated cells (Figure 4C), we probed its transcriptional regulation by FoxO3a transcription factor whose phosphorylation leads to its inactivation and sequestration in the cytoplasm while its non phosphorylated form translocates to the nucleus and transcribes the p27 gene to induce G1 arrest (49, 51, 63, 65, 66). We found that DDR2 knockdown in mitogen-stimulated cells reduces FoxO3a phosphorylation (Figure 6A), leading to its activation and nuclear localization (Figure 6B), which corresponded with enhanced p27 and reduced PCNA levels (Figure 4C).

**Figure 6:**
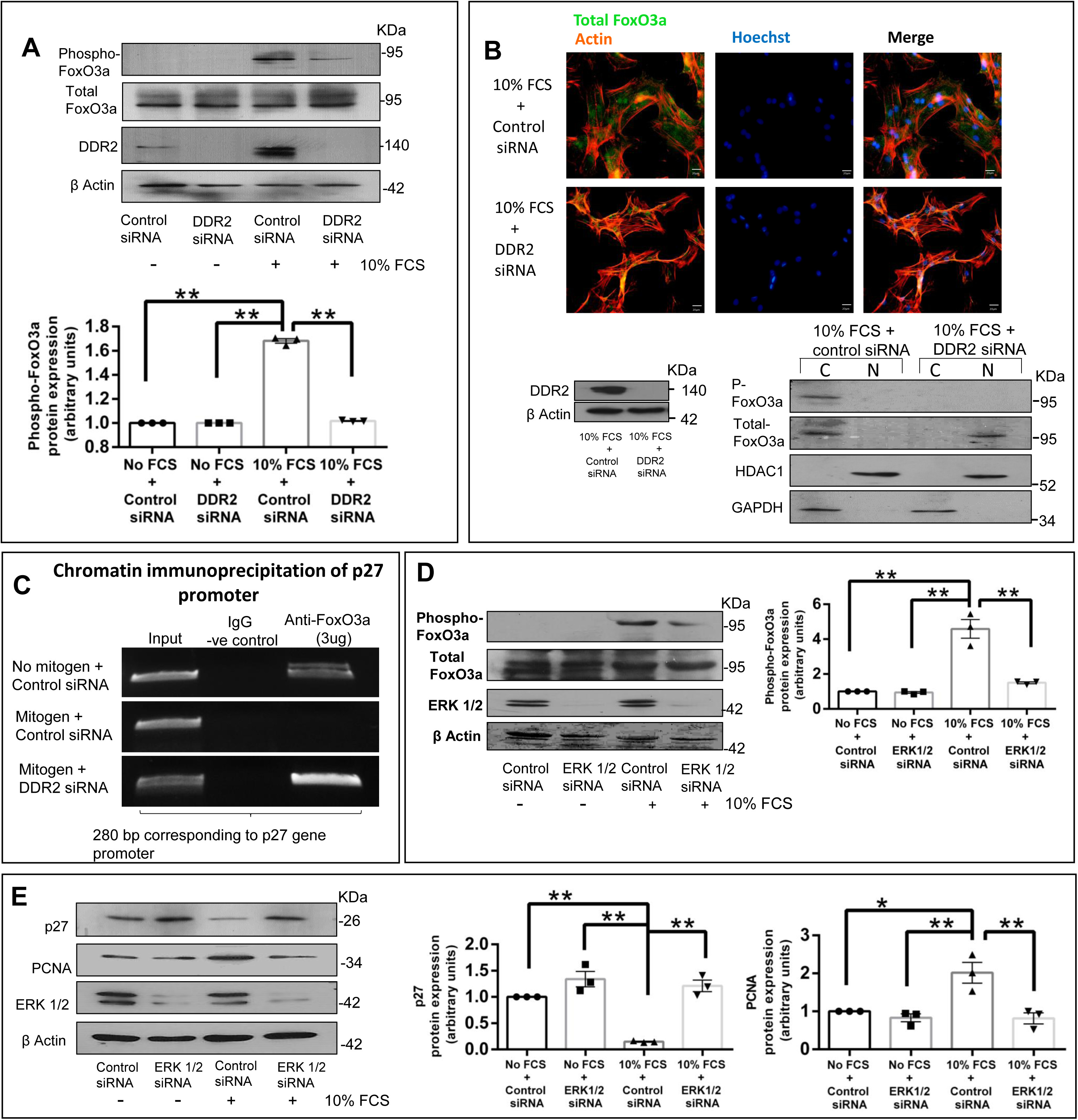
Regulation of p27 by DDR2: ii) Transcriptional regulation through modulating FoxO3a activity. (**A-E**) Sub-confluent cultures of cardiac fibroblasts were transfected with DDR2 siRNA (**A, B and C**) or ERK1/2 siRNA (**D and E**) (with control siRNA) and, following revival for 12h in 10% serum-supplemented medium, the cells were serum-deprived for synchronization. Post-synchronization, the cells were exposed to 10% Fetal Calf Serum. (**A**) DDR2 siRNA-transfected cells were collected at 8h and analysed by western blotting for expression levels of Phospho-FoxO3a (T32)/Total FoxO3a, with β-actin as loading control. Significance was determined by two-way ANOVA (Tukey’s multiple comparisons test, **p< 0.01, comparisons as depicted in the Figure). Validation of DDR2 knockdown is also shown. (**B**) Immunocytochemistry for Total FoxO3a/actin (upper colour panel) and nuclear-cytoplasmic isolation (lower panel) was performed on cells 8h after addition of 10% FCS, according to the protocol under Methods. The Figure shows (Colour panel) Total Foxo3a stained by Alexa 488 secondary antibody (Green) and Actin counterstained by phalloidin-rhodamine (Red). Nucleus is stained with Hoechst (Blue). DDR2 siRNA-treated cells show all nuclear staining for FoxO3a. The lower panel shows the western blot analysis of cytoplasmic (C) and nuclear (N) fractions isolated following treatments. Phospho-FoxO3a and Total Foxo3a were analysed along with HDAC1 as marker for nuclear fraction and GAPDH for cytoplasmic fraction. Validation of DDR2 knockdown from the same set of experiment is shown (lower left panel). (**C**) DDR2 siRNA-transfected cells were collected at 30 min and chromatin was immunoprecipitated using anti-FoxO3a antibody followed by PCR amplification and analysed on a 2% agarose gel for presence of the 280bp region of the p27 gene promoter (region amplified is specified in methods section). (**D**) ERK1/2 MAPK siRNA-transfected cells were collected at 8h and analysed by western blotting for expression levels of Phospho-FoxO3a (T32)/total FoxO3a, with β-actin as loading control. Significance was determined by two-way ANOVA (Tukey’s multiple comparisons test, **p< 0.01, comparisons as depicted in the Figure). Validation of ERK1/2 MAPK knockdown is also shown. (**E**) ERK1/2 MAPK siRNA-transfected cells were collected at 8h and analysed by western blotting for expression levels of p27 and PCNA, with β-actin as loading control. Significance was determined by two-way ANOVA (Tukey’s multiple comparisons test, *p<0.05 and **p<0.01, comparisons as depicted in the Figure). Data are representative of 3 independent experiments, n=3. Mean ± SEM (Standard Error of Mean).

Consistent with these observations, chromatin immunoprecipitation showed enhanced binding of FoxO3a to the p27 gene promoter in DDR2-silenced, mitogen-stimulated cells (Figure 6C).

Subsequently, we also examined the mechanism by which DDR2 regulates FoxO3a. While DDR2 knockdown inhibited ERK1/2 MAPK in serum-stimulated cells (Figure 5D), ERK1/2 MAPK knockdown in mitogen-stimulated cells reduced FoxO3a phosphorylation (Figure 6D), resulting in enhanced p27 and reduced PCNA (Figure 6E).The data show that DDR2 promotes ERK1/2 MAPK activation to inhibit FoxO3a-dependent transcriptional induction of p27, promoting G1-S transition.

Considered in tandem, the data point to SRF/Skp2-dependent post-translational and FoxO3a dependent transcriptional regulation of p27 by DDR2.

#### DDR2 over-expression facilitates G1-S transition in mitogen-deprived cardiac fibroblasts

Cardiac fibroblasts were transfected with a plasmid construct containing DDR2 cDNA driven by a CMV promoter and grown under serum-free conditions. Over-expression of DDR2 in mitogen-starved cells promoted G1-S transition, demonstrated by flow cytometry (Figure 7A). This was accompanied by enhanced Skp2 levels, reduced p27 levels, elevated cyclin D1/E levels, enhanced Rb phosphorylation and enhanced PCNA expression (Figures 7B and C). Further, DDR2 over-expression also led to ERK1/2 MAPK activation (Figure 7D), a significant increase in SRF levels (Figure 7E) and elevated Phospho-FoxO3a (inactivation) levels (Figure 7E), resulting in its cytoplasmic localisation (Figure7F). As expected, ERK1/2 MAPK inhibition using siRNA or PD98059 in DDR2-over-expressing cells attenuated PCNA (Figure 7G) and Phospho-FoxO3a levels (indicating FoxO3a activation), and enhanced p27 expression (Figure 7H), showing that ERK1/2 MAPK mediates the DDR2 effects.

**Figure 7:**
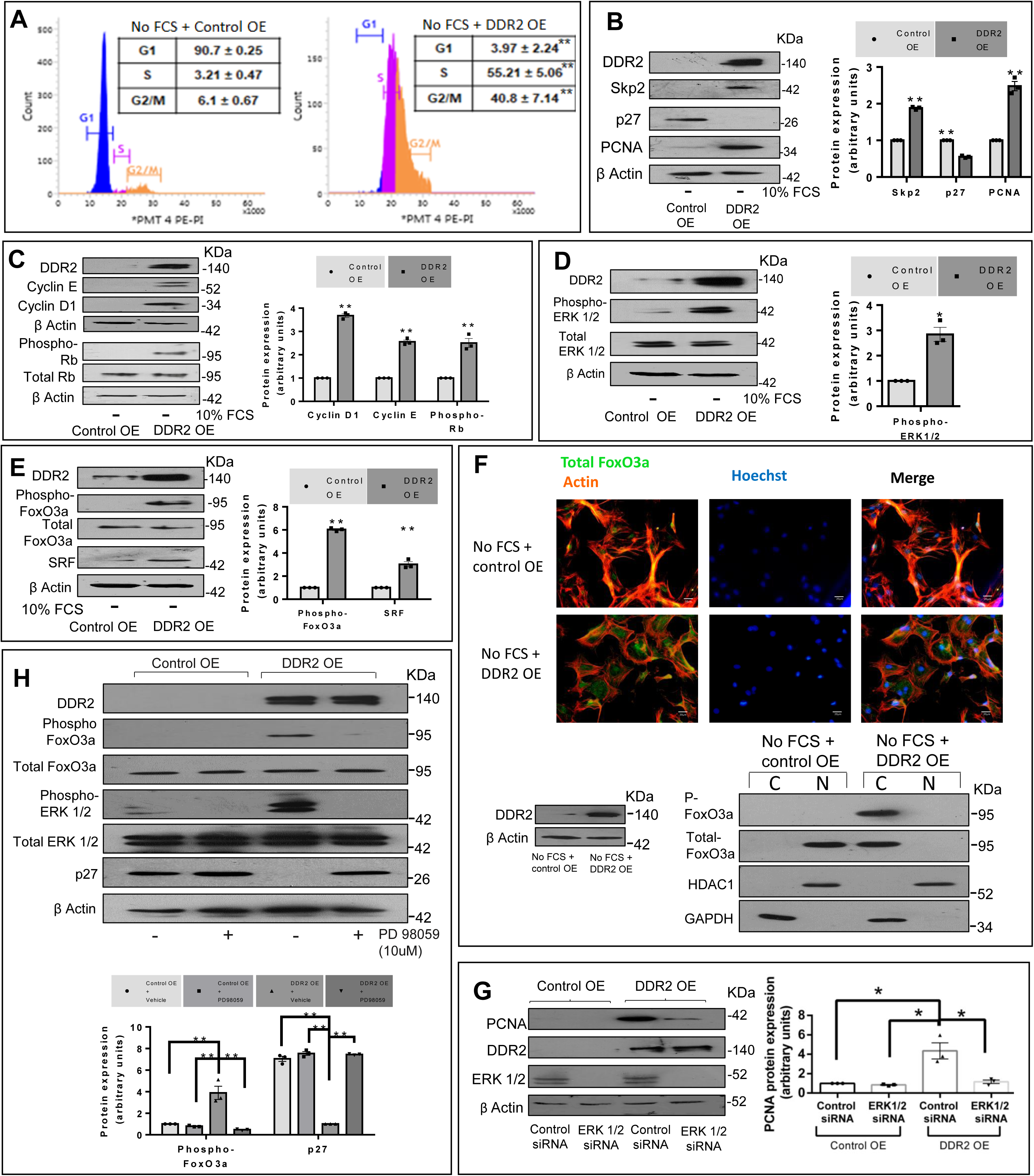
DDR2 over-expression facilitates G1-S transition in mitogen-deprived cardiac fibroblasts. (**A-E**) Cardiac fibroblasts transfected with DDR2 cDNA over-expression plasmid (DDR2 OE) or empty vector control (Control OE) were subjected to western blot analysis (with β-actin as loading control) or flow cytometric analysis. Validation of DDR2 overexpression is also shown. (**A**) Flow cytometric profile of G1-S transition, showing distribution of cells in each phase. Significance was determined by Student’s t test, **p< 0.01 vs Control OE. (**B**) Skp2, p27 and PCNA protein levels were examined. Significance was determined by Student’s t test, **p< 0.01vs Control OE. (**C**) Cyclin E, Cyclin D1 and Phospho-Rb/Total Rb protein levels were examined by western blot analysis. Significance was determined by Student’s t test, **p< 0.01 vs Control OE. (**D**) Phospho-ERK1/2 / total ERK1/2 MAPK protein level. Significance was determined by Student’s t test, *p< 0.05 vs Control OE. (**E**) Phospho-FoxO3a/Total FoxO3a and SRF protein level. Significance was determined by Student’s t test, **p< 0.01 vs Control OE. (**F**) Cardiac fibroblasts transfected with DDR2 cDNA over-expression plasmid (DDR2 OE) or empty vector control (Control OE) were grown in serum-free medium for 24h and subjected to immunocytochemistry (colour panel) for Total Foxo3a (Green) counterstained with actin (Phalloidin-rhodamine, Red) and nuclear staining by Hoechst (Blue). In DDR2 OE cells, Total FoxO3a is stained in the cytoplasmic region and, in Control OE cells, Total-FoxO3a stained in the nucleus. (Lower panel) Nuclear-cytoplasmic isolation and analysis by western blotting for Phospho-FoxO3a and Total FoxO3a along with nuclear fraction marker, HDAC1, and cytoplasmic marker, GAPDH. In DDR2 OE cells, FoxO3a shows phosphorylation and is co-localised with GAPDH (cytoplasmic marker) and in Control OE cells, FoxO3a is not phosphorylated and is found along with HDAC1 (nuclear marker). Validation of DDR2 over-expression from the same set is shown (lower left panel). (**G**) Cardiac fibroblasts were co-transfected with DDR2 cDNA over-expression plasmid (DDR2 OE) with empty vector control (Control OE) and ERK1/2 MAPK siRNA (with Control siRNA), post-revival, the cells were serum-deprived for 24h and subjected to western blot analysis for protein levels of PCNA (with β-actin as loading control). Significance was determined by two-way ANOVA (Tukey’s multiple comparisons test, *p<0.05, comparisons as depicted in the Figure). DDR2 over-expression validation and validation of ERK1/2 MAPK knockdown are also shown. (**H**) Cardiac fibroblasts were transfected with DDR2 cDNA over-expression plasmid (DDR2 OE) with empty vector control (Control OE). Following 24h of transfection, the cells were treated with PD 98059 (10μM) (ERK1/2 inhibitor) for 8h and then analysed by western blotting for Phospho-FoxO3a levels and p27 levels. (with Total FoxO3a and β-actin as loading controls, respectively). Significance was determined by two-way ANOVA (Tukey’s multiple comparisons test, **p< 0.01, comparisons as depicted in the Figure). Data are representative of 3 independent experiments, n=3. Mean ± SEM (Standard Error of Mean).

#### Enhanced expression of DDR2 correlates with enhanced levels of SRF, cIAP2 and PCNA in freshly isolated cardiac fibroblasts from Spontaneously Hypertensive Rats

Looking for a possible association of DDR2 with augmented cIAP2 expression and cardiac fibroblast hyperplasia in Spontaneously Hypertensive Rats (SHR), we analysed expression levels of DDR2, SRF, cIAP2 and PCNA in freshly prepared cardiac fibroblasts (following 2.5h of pre-plating of freshly isolated cells) from 6-month old male SHR. Consistent with the in vitro findings, enhanced levels of DDR2 in cardiac fibroblasts from SHR correlated well with enhanced levels of SRF, cIAP2 and PCNA (Figure 8A). Further, while serum from both SHR and control rats enhanced the levels of DDR2, SRF, cIAP2 and PCNA in cardiac fibroblasts in vitro, serum from SHR had more pronounced effects (Figure 8B), which were reduced upon pre-treatment of the cells with candesartan.

**Figure 8:**
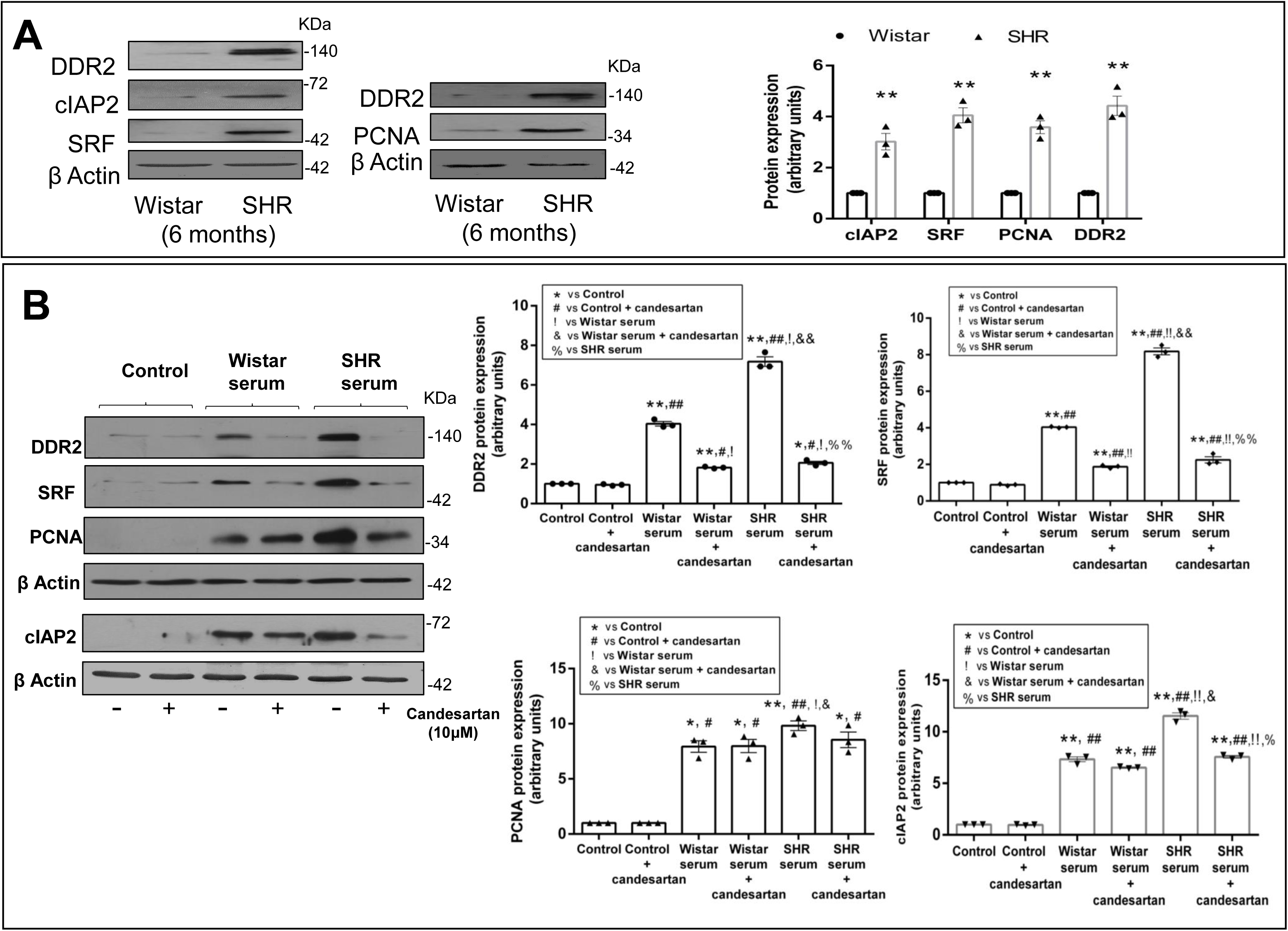
Enhanced expression of DDR2 correlates with enhanced levels of SRF, cIAP2 and PCNA in freshly isolated cardiac fibroblasts from Spontaneously Hypertensive Rats. (**A**) Cardiac fibroblasts freshly isolated from 6-month old male SHR and Wistar rats were analysed by western blotting for levels of cIAP2, SRF, PCNA and DDR2 with β-actin as loading control. Significance was determined by Student’s t test, ** p< 0.01 vs Wistar. (**B**) Synchronized sub-confluent cultures of cardiac fibroblasts from Wistar rats were exposed to serum obtained from male SHR or Wistar rats (6-months) with or without pre-incubation with 10μM candesartan. After 12h incubation, DDR2, SRF, PCNA and cIAP2 protein levels were examined in these cells, with β-actin as loading control. Significance was determined by two-way ANOVA (Tukey’s multiple comparisons test, *, #, !, & and % p<0.05 vs Control, Control + candesartan, Wistar serum, Wistar serum + candesartan and SHR serum, respectively (as depicted in the Figure). Double symbols represent p< 0.01. Data are representative of 3 independent experiments, n=3. Mean ± SEM (Standard Error of Mean).

##### Obligate role of collagen type I in the activation of DDR2 and its regulatory role in cell survival and cell cycle protein expression

Treatment of cardiac fibroblasts with Ang II, H_2_O_2_ and 10% FCS caused activation of DDR2, as shown by enhanced Phospho-tyrosine levels in immunoprecipitated DDR2, which was attenuated by WRG-28, a specific inhibitor of collagen type I binding to DDR2 (16), indicating activation of DDR2 by collagen type I (Figures 9A-D). Further, activation of DDR2 in response to serum from SHR was also inhibited by WRG-28 (Figure 9E). Notably, WRG-28 treatment attenuated the expression of cIAP2, Skp2, PCNA, p27, SRF, Phospho-ERK1/2 and Phospho-FoxO3a levels (Figures 10 A-E), which was similar to the effects of DDR2 knockdown. Moreover, apoptosis was evident in WRG-28-treated cells exposed to oxidative stress, as shown by an increase in cleaved-caspase-3 level (Figure 10 F). A schematic representation of the plausible molecular events that integrate apoptosis resistance and proliferation under the regulatory control of DDR2 in cardiac fibroblasts is provided in Figure 11.

**Figure 9:**
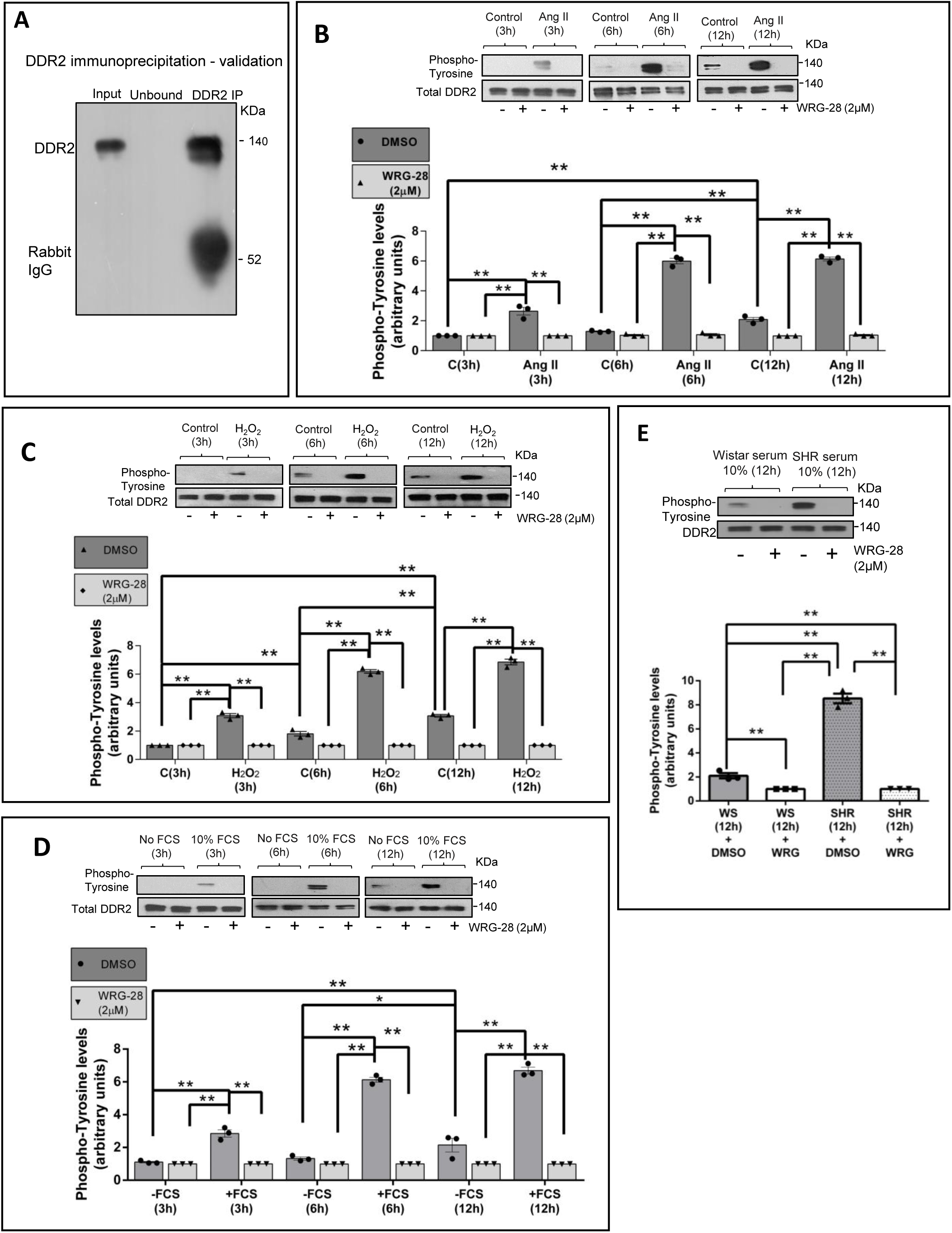
Obligate role of collagen type I in the activation (phosphorylation) of DDR2. (**A**). Validation blot for immunoprecipitation of DDR2. DDR2 was pulled down from cardiac fibroblast cell lysates using specific DDR2 anti-rabbit IgG (2ug antibody for 100ug of total cell lysate) and protein A magnetic beads. Western blot was performed to confirm the pull down of DDR2. In the blot, “input” indicates the fraction of total cell lysate before immunoprecipitation, and “unbound” the supernatant of the immunoprecipitated sample. Lane 3 is the immunoprecipitated DDR2 (DDR2 IP), showing DDR2 pull down. (B-D) Following serum deprivation for 24h, cardiac fibroblasts were divided into 4 groups and treated as follows: 1) Control + WRG-28 (2μM), 2) Control + DMSO, 3) Stimulus + WRG-28 (2μM), and 4) Stimulus + DMSO. Ang II (1μM) or H_2_O_2_ (25μM) or 10% Fetal Calf Serum (10 %FCS) was used as the stimulus. Before treatment with the stimulus, the cells were pre-incubated with either WRG-28 (2μM) or vehicle DMSO for 45 min. Following preincubation, the cells were treated with the stimulus for 3, 6 and 12h and total DDR2 was immunoprecipitated (see also supplementary figure S2A) and subjected to western blot analysis of Phospho-tyrosine levels (activated DDR2) normalized to total DDR2 levels. (**B**) Phospho-tyrosine levels in immunoprecipitated DDR2 at 3, 6 and 12h of Ang II (1μM) treatment. Significance was determined by two-way ANOVA (Tukey’s multiple comparisons test, **p<0.01, comparisons as depicted in the figure, ANOVA summary: p <0.01 for Interaction (stimulus), Row Factor (hours) and Column Factor (WRG-28)) (**C**) Phosphotyrosine levels in immunoprecipitated DDR2 at 3, 6 and 12h of H_2_O_2_ (25μM) treatment. Significance was determined by two-way ANOVA (Tukey’s multiple comparisons test, **p<0.01, comparisons as depicted in the figure, ANOVA summary: p <0.01 for Interaction (stimulus), Row Factor (hours) and Column Factor (WRG-28)) (**D**) Phospho-tyrosine levels in immunoprecipitated DDR2 at 3, 6 and 12h of 10% Fetal Calf Serum(10% FCS) treatment. Significance was determined by two-way ANOVA (Tukey’s multiple comparisons test, **p<0.01, comparisons as depicted in the figure, ANOVA summary: p <0.01 for Interaction (stimulus), Row Factor (hours) and Column Factor (WRG-28)) (**E**) Cardiac fibroblasts were treated with serum isolated from Wistar rats or SHR following WRG-28 or DMSO pretreatment for 45 min. After 12h, DDR2 was immunoprecipitated and analysed for phosphotyrosine levels, with total DDR2 as loading control. Significance was determined by two-way ANOVA (Tukey’s multiple comparisons test, **p<0.01, comparisons as depicted in the figure). Data are representative of 3 independent experiments, n=3. Mean ± SEM (Standard Error of Mean).

**Figure 10:**
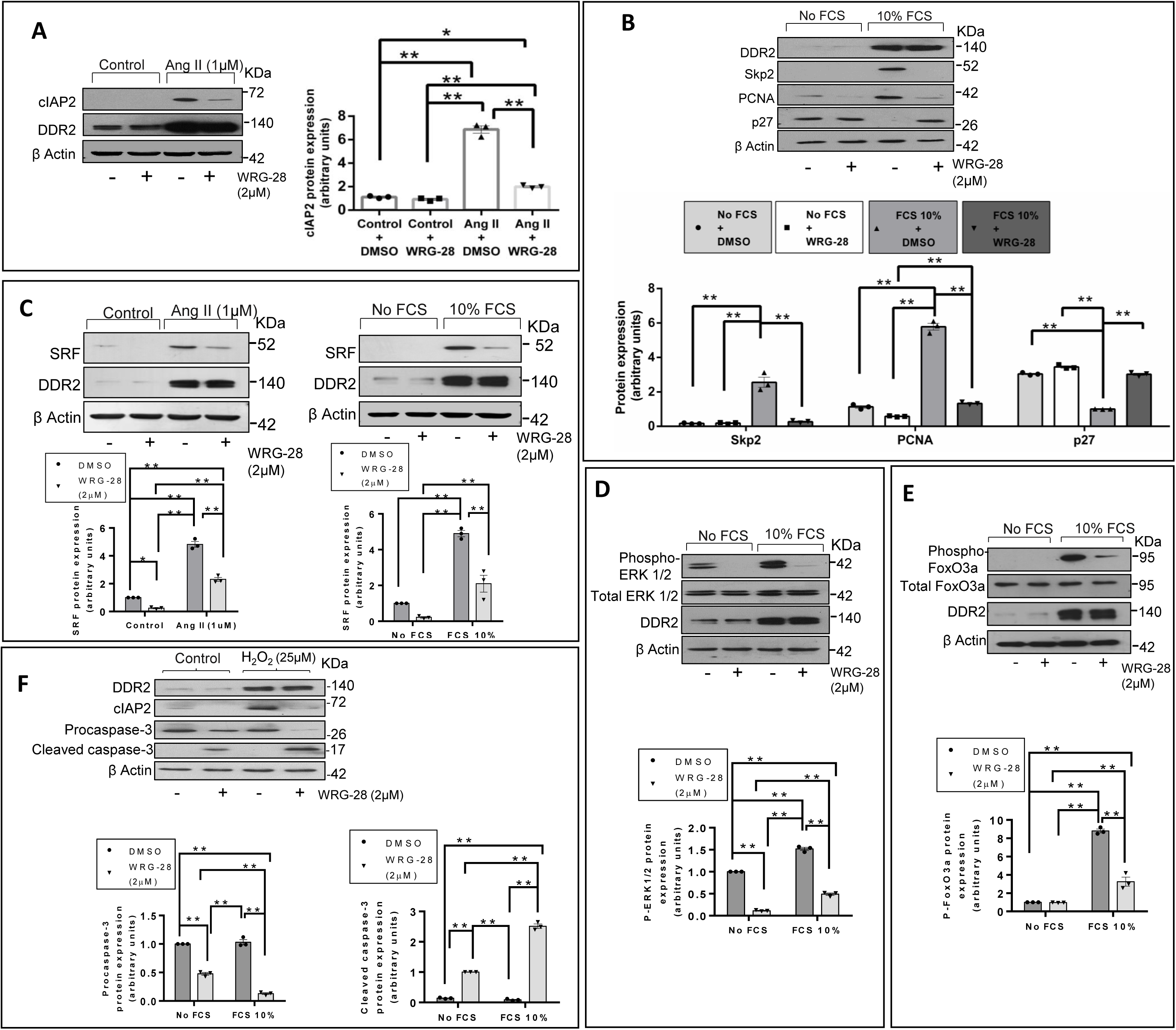
Obligate role of collagen type I in the regulation of cell survival and cell cycle protein expression. (**A**) Cardiac fibroblasts were treated with Ang II (1μM) following pre-treatment with WRG-28 (2μM) or DMSO for 45 min. Cells were collected after 12h of Ang II treatment and analysed by western blotting for expression levels of cIAP2, with β-actin as loading control. Significance was determined by two-way ANOVA (Tukey’s multiple comparisons test, *p<0.05 and **p<0.01, comparisons as depicted in the figure). (**B**) After 24h of serum deprivation, cardiac fibroblasts were pre-treated for 45 min with WRG-28 (2μM) or DMSO, then exposed to 10% Fetal Calf Serum (10% FCS) for 8h and analysed by western blotting for expression levels of Skp2, PCNA and p27, with β-actin as loading control. Significance was determined by two-way ANOVA (Tukey’s multiple comparisons test, **p<0.01, comparisons as depicted in the Figure). (**C**) Cardiac fibroblasts were treated for 12 h with Ang II (1μM) or 10% FCS after pre-treatment for 45 min with WRG-28 (2μM) or DMSO, and analysed by western blotting for expression levels of SRF, with β-actin as loading control. Significance was determined by two-way ANOVA (Tukey’s multiple comparisons test, **p<0.01, comparisons as depicted in the figure). (**D**) Following serum deprivation for 24h, cardiac fibroblasts were treated with 10% FCS for 8h after pre-treatment for 45 min with WRG-28 (2μM) or DMSO and analysed by western blotting for expression levels of Phospho-ERK1/2, with Total ERK1/2 as loading control. Significance was determined by two-way ANOVA (Tukey’s multiple comparisons test, **p<0.01, comparisons as depicted in the figure). (**E**) Following serum deprivation for 24h, cardiac fibroblasts were treated with 10% FCS for 8h after pre-treatment for 45 min with WRG-28 (2μM) or DMSO and analysed by western blotting for expression levels of Phospho-FoxO3a, with Total FoxO3a as loading control. Significance was determined by two-way ANOVA (Tukey’s multiple comparisons test, **p<0.01, comparisons as depicted in the figure). (**F**) Following pre-treatment for 45 min with WRG-28 (2μM) or DMSO and treatment with H_2_O_2_ (25μM) for 8h, cells were collected and analysed by western blot for the expression levels of cleaved-caspase 3 and procaspase 3, with β-actin as loading control. Significance was determined by two-way ANOVA (Tukey’s multiple comparisons test, **p<0.01, comparisons as depicted in the figure). Data are representative of 3 independent experiments, n=3. Mean ± SEM (Standard Error of Mean).

**Figure 11:**
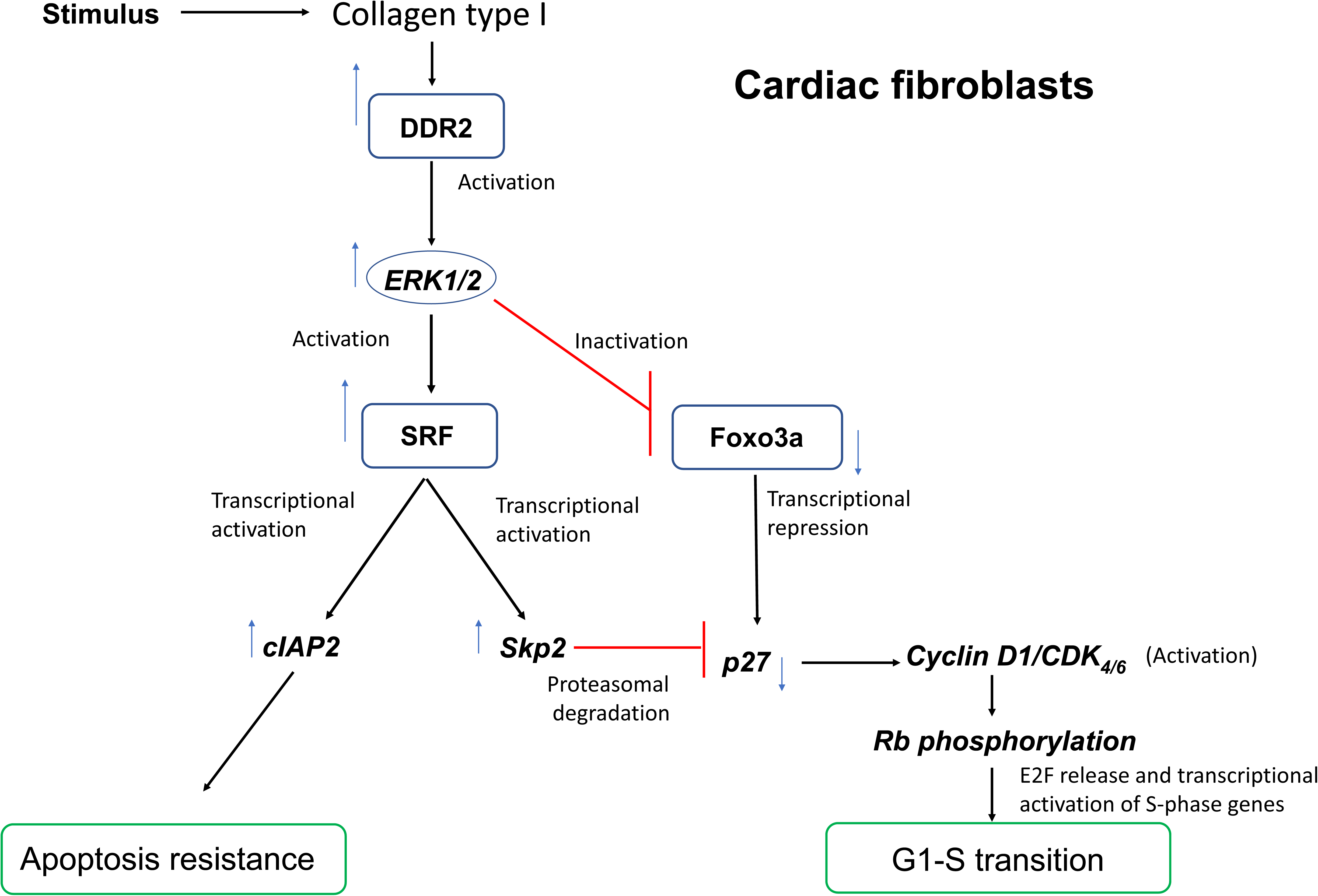
A schematic representation of the plausible molecular events that integrate apoptosis resistance and proliferation under the regulatory control of DDR2 in cardiac fibroblasts. In response to the stimulus (Ang II or 10% FCS), collagen type I-dependent activation of DDR2 leads to the activation of ERK1/2 MAPK that in turn activates SRF to: i) transcriptionally increase cIAP2, conferring apoptosis resistance, ii) increase Skp2 and promote Skp2-mediated *post-translational* degradation of p27, and iii) inactivate FoxO3a, through phosphorylation, to transcriptionally repress p27. Transcriptional and posttranslational inhibition of p27 results in Cyclin D1/CDK4/6 complex-dependent phosphorylation of Rb protein, facilitating the transcription of S-phase genes and G1-S transition.

## Discussion

Cardiac fibroblasts, an abundant cell type in the heart, are the only intracardiac source of collagen types I and III. Although their ability to survive and proliferate in the hostile ambience of the injured myocardium and generate a fibrillar collagen-rich scar underlies wound healing in the short-term, their persistence in the infarct scar due to relative resistance to apoptosis promotes adverse myocardial remodeling in the long-term, which exacerbates ventricular dysfunction and contributes to the progression of heart failure. A better understanding of the molecular pathways that regulate apoptosis resistance and cell cycle progression in cardiac fibroblasts can contribute to our ability to limit adverse fibrotic remodeling of the heart.

After examining several apoptotic regulators, a role for constitutively-expressed Bcl2 in protecting cardiac fibroblasts against a variety of pro-apoptotic stimuli has been reported earlier (38). On the other hand, we had demonstrated that augmented cIAP2 expression protects cardiac fibroblasts against oxidative damage (48). The present study sought to analyze the role of DDR2 as a possible pro-survival factor in cardiac fibroblasts, focusing on its regulatory role in cIAP2 expression. Currently, there are only sporadic studies that report the anti-apoptotic role of DDR2 in tumour cell lines and cell types such as pulmonary fibroblast and hepatic stellate cells, with limited mechanistic insights (8, 23, 31, 34). We provide robust evidence that Ang II enhances cIAP2 expression via DDR2-dependent activation of ERK1/2 MAPK that in turn leads to SRF activation and binding to the cIAP2 gene promoter (Figures 1 and 2). Additionally, the involvement of MRTF-A/B, a cofactor of SRF (45, 62), in the cellular response to Ang II was evident since its inhibition led to down regulation of cIAP2 protein expression (Figure 2A). It is to be noted that CCG-1423 inhibits the MRTF-A and MRTF-B isoforms (20), both of which are expressed in fibroblasts (7, 47, 62). It is pertinent to point out here that both Ang II and ERK1/2 MAPK have been shown to activate SRF (9, 19, 33, 56). Further, SRF has been identified as an important factor in myofibroblast differentiation leading to tissue fibrosis (4, 54, 67), and is reported to play a role in cell survival and cell proliferation in other cell types (52, 64). Exploring the significance of Ang II-dependent increase in cIAP2 expression, we found that exposure of cells to oxidative stress (25μM H_2_O_2_) enhances DDR2, SRF and cIAP2 levels while knockdown of DDR2 or SRF abolishes cIAP2 expression and causes cell death under conditions of oxidative stress (Figures 3A and C-G). Interestingly, candesartan, the AT1 receptor antagonist, attenuated H_2_O_2_-stimulated DDR2, SRF and cIAP2 expression and compromised viability in cardiac fibroblasts (Figures 3A and B), showing that oxidative stress induces Ang II production that in turn protects the cells against oxidative injury via DDR2-dependent cIAP2 induction. Notably, over-expression of cIAP2 gene was found to abolish the activation of apoptosis in DDR2-silenced cardiac fibroblasts exposed to H_2_O_2_, underscoring the centrality of cIAP2 in protecting the cells (Figure 3H). It is also noteworthy that serum from SHR had a stimulatory effect on DDR2, SRF and cIAP2 expression in cardiac fibroblasts isolated from normal rats, which was reduced by candesartan (Figure 8B), implying a role for circulating Ang II in SHR in triggering pro-survival mechanisms in cardiac fibroblasts.

These findings need to be considered in tandem with earlier reports that Ang II induces apoptosis in cardiac myocytes (2, 24). The pro-apoptotic effect of Ang II on cardiac myocytes, on the one hand, and its anti-apoptotic effect on cardiac fibroblasts, on the other, constitute yet another example of “the apoptotic paradox” that was previously reported in a setting of idiopathic pulmonary fibrosis wherein TGF-β1 exerts pro-apoptotic effects on epithelial cells and anti-apoptotic effects on myofibroblasts in the lung (57). The pro-survival role of Ang II alongside its stimulatory effect on collagen expression in cardiac fibroblasts may facilitate fibroblast-mediated wound healing upon injury in the short-term but may also contribute to the persistence of these cells in the scar in an active state long after the termination of the healing process, which ultimately would lead to myocardial stromal expansion and pump dysfunction. Ang II is recognized to be a potent pro-fibrotic factor and its role in cardiac fibroblast function has been viewed mostly in terms of its effects on collagen turnover (55). By focusing on the anti-apoptotic action of Ang II on cardiac fibroblasts, mediated by DDR2 signaling, this study uncovers a less known facet of the pro fibrotic action of Ang II and, possibly, offers an additional explanation for the beneficial effects of Ang II inhibitors that are used extensively in the clinical setting and are known to minimize adverse myocardial remodeling post myocardial injury (18, 40, 50, 53).

Beyond demonstrating the obligate role of DDR2 in cell survival, this study also uncovers a critical role for DDR2 in cell cycle progression in cardiac fibroblasts. While a role for DDR2 in promoting proliferation in hepatic stellate cells, skin fibroblasts, chondrocytes and cancer cell types has been demonstrated (17, 28, 36, 42, 43), an anti-proliferative and lack of proliferative effect of DDR2 have also been reported in other cell types (25, 41, 61). Moreover, these studies did not probe the direct influence of DDR2 on cell cycle regulatory elements. Against this backdrop, the present study provides evidence of the direct involvement of DDR2 in the cell cycle machinery to facilitate G1-S transition in mitogen stimulated cardiac fibroblasts.

G1-S transition occurs following activation of cyclin-dependent kinases (CDKs) through association with cyclins, and consequent phosphorylation of the retinoblastoma protein (Rb) and E2F-dependent transcription of S-phase genes, including PCNA and Cyclin E. On the other hand, inhibition of CDK activity by cyclin-dependent kinase inhibitors (CDKIs) leads to Rb hypophosphorylation and cell cycle arrest (21, 37). The abundance of p27Kip1, an important member of the Cip/Kip family of CDKIs and critical regulator of G1-S transition, is regulated by post-translational and transcriptional mechanisms. S-phase kinase-associated protein 2 (Skp2), an F-box protein of the SCF ubiquitin ligase complex (5, 11), facilitates cell cycle progression through the proteasomal degradation of p27. p27 is also regulated transcriptionally (26, 29). Forkhead box O 3a (FoxO3a), a member of the Forkhead box O (FoxO) family of transcription factors, enhances p27 gene expression, and phosphorylation dependent inhibition of FoxO3a activity facilitates cell-cycle progression through transcriptional repression of p27 in mitogen-stimulated cells (49, 51, 63, 65, 66). This study focused on the regulation of these mediators by DDR2 and sought to delineate the underlying mechanisms.

Flow cytometry demonstrated that DDR2 knockdown in mitogen-stimulated cells causes cell cycle arrest at the G1 phase (Figure 4B), accompanied by a significant reduction in the levels of PCNA, along with reduced Skp2 levels, induction of p27 (Figure 4C), and hypophosphorylation of Rb (Figure 4D). Further, cyclin E but not cyclin D1 was reduced in DDR2-silenced, mitogen-stimulated cells (Figure 4E). Cyclin D1 serves as a key sensor and integrator of extracellular signals of cells in early to mid-G1 phase whereas cyclin E is synthesized during progression to S phase and it associates with Cdk2 and activates its kinase activity shortly before entry of cells into the S phase (39). Thus, our data suggest that DDR2 may act at the G1 phase, preventing S phase entry by inhibiting Cyclin D1/CDK activation through p27 induction. Additional evidence of an obligate role for DDR2 in G1-S transition came from experiments showing that over-expression of DDR2 in mitogen-deprived cells increases the expression of cyclin D1, cyclin E and Skp2, reduced p27 and promoted Rb phosphorylation, culminating in enhanced PCNA expression and G1-S transition (Figures 7A-C). It is noteworthy that, while DDR2 over-expression in mitogen-starved cells induced cyclin D1, DDR2 knockdown in mitogen-stimulated cells did not inhibit cyclin D1 expression, indicating that serum may overcome the effect of DDR2 knockdown, a possibility that warrants further investigation.

Focusing on the mechanisms involved in cell cycle regulation by DDR2, we found that DDR2 facilitates G1-S transition through FoxO3a-mediated transcriptional and SRF/Skp2-mediated post-translational inhibition of p27 expression. Our findings showed that DDR2 knockdown in mitogen-stimulated cells leads to inhibition of ERK1/2 activity (Figure 5D) and consequent FoxO3a activation and nuclear translocation through its hypophosphorylation (Figure 6A,B and D), which in turn transcriptionally enhances p27 expression, as shown by FoxO3a binding to the p27 gene promoter (Figure 6C). On the other hand, DDR2 over-expression in mitogen-starved cells led to enhanced ERK1/2 activity (Figure 7D) and hyperphosphorylation of FoxO3a and its cytoplasmic localisation (Figures 7E and F). However, inhibition of ERK1/2 activation in DDR2-over-expressed cells resulted in hypophosphorylation of FoxO3a (activation) and increased p27 expression (Figure 7H), consistent with down-regulation of PCNA levels in ERK1/2-silenced DDR2 over-expressing cells (Figure 7G). Thus, DDR2 acts as a negative regulator of FoxO3a. Further, while DDR2 knockdown in mitogen-stimulated cells resulted in ERK1/2 MAPK inhibition (Figure 5D) and consequent down-regulation of SRF (Figure 5B and C), SRF knockdown reduced Skp2 levels (Figure 5A), resulting in increased p27, reduced PCNA and G1-S transition (Figures 5F and G). Together with the observation that over-expression of DDR2 in mitogen-deprived cells increases the expression of SRF and Skp2 (Figures 7B and C), these data clearly show that DDR2 is a positive regulator of Skp2, as confirmed by SRF binding to the Skp2 gene promoter, which was abolished upon DDR2 silencing (Figure 5E). To the best of our knowledge, there is only a single report showing the link between SRF and Skp2 expression (64). Together, the data demonstrate that DDR2-induced ERK1/2 MAPK activation promotes transcriptional repression of p27 through FoxO3a phosphorylation and SRF-mediated post translational degradation of p27 by Skp2.

It is pertinent to note that, apart from elucidating the obligate role of DDR2 in cell survival and cell cycle progression in cardiac fibroblasts, this study uncovers a common pathway of regulation of these processes wherein the DDR2-ERK1/2 MAPK-SRF signaling pathway regulates apoptosis resistance via cIAP2 and cell cycle progression via Skp2 that promotes proteasomal degradation of p27 to facilitate Rb phosphorylation and G1-S transition. Notably, the findings also point to the obligate role of collagen type I in the activation of DDR2 (Figures 9A-E) and its regulatory role in cell survival and cell cycle (Figures 10 A-F). The role of DDR2 as an important determinant of cardiac size and organ growth was previously demonstrated in DDR2-null mice (6). Investigations on this mouse model had demonstrated, by echocardiography, reduced left ventricular chamber dimensions and reduced cardiomyocyte length, resulting in decreased heart size and weight. Moreover, the study reported a reduction in cardiac interstitial collagen density at baseline and reduced rates of collagen synthesis in cardiac fibroblasts isolated from DDR2-null mice, compared to the wild type littermate fibroblasts. In the present study, the link between DDR2 and factors that mediate apoptosis resistance and G1-transition in an in vivo setting of hypertension-induced myocardial disease underscored the role of DDR2 in regulating cardiac fibroblast growth. We found significantly increased levels of DDR2 that correlated well with increased levels of SRF, cIAP2 and PCNA in cardiac fibroblasts freshly isolated from Spontaneously Hypertensive Rats (Figure 8A), which present a genetic model of hypertensive heart disease with a role for the renin-angiotensin system in mediating the pathological changes in the myocardium **(15)**. Interestingly, serum from SHR induced the expression of DDR2, SRF and PCNA in cardiac fibroblasts from normal rats, which was attenuated by candesartan (Figure 8B), indicating the effect of circulating Ang II on cardiac fibroblasts in SHR rats.

In summary, cardiac fibroblasts are major contributors to tissue repair following acute cardiac injury, and to cardiac fibrogenesis associated with most forms of chronic heart disease. Phenotypic transformation, proliferation, collagen production and persistence *post healing* due to apoptosis resistance are distinct attributes of cardiac fibroblasts that are key to their pivotal role in myocardial pathophysiology. Identification of cardiac fibroblast-specific factors and their downstream effectors that regulate these processes is therefore of considerable scientific interest with obvious clinical relevance. Our earlier studies had demonstrated an obligate role for DDR2 in collagen expression in cardiac fibroblasts (14) and in their phenotypic transformation in response to Ang II (59). In fact, our recent investigations have identified DDR2 as a possible molecular link between arterial fibrosis and metabolic syndrome in rhesus monkeys (58), with an obligate role in the regulation of collagen gene expression in vascular cells as well. In the present communication, we present evidence that DDR2-dependent ERK1/2 MAPK activation facilitates the coordinated regulation of apoptosis resistance and cell cycle progression in cardiac fibroblasts via SRF. Together, the data place these cardinal aspects of cardiac fibroblast function within a single mechanistic framework of molecular pathways under the regulatory control of DDR2, which defines the fibroblast phenotype and acts as a ‘master switch’ in these cells. The predominant localization of DDR2 in cardiac fibroblasts and its regulatory role in cardiac fibroblast function lend support to the postulation (14) that it is a potential drug target in the control of cardiac fibroblast-mediated adverse tissue remodeling.

## Abbreviations

DDR2: Discoidin Domain Receptor 2
SRF: Serum Response Factor
ERK1/2 MAPK: Extracellular signal-regulated kinase1/2 Mitogen-activated Protein Kinase
cIAP2: Cellular inhibitors of apoptosis protein 2
FoxO3a: Forkhead box O 3a transcription factor
Skp2: S-Phase Kinase Associated Protein 2

## Acknowledgements

This work was supported by a research grant to SK from the Department of Biotechnology, Government of India (BT/PR23486/BRB/10/1589/2017). AST thanks SCTIMST for a Research Fellowship. HV thanks the Department of Biotechnology, Government of India, for a Research Fellowship. The authors thank Dr Rakesh Laishram of the Rajiv Gandhi Centre for Biotechnology, Trivandrum, for providing Phospho-tyrosine antibody and Dr Lakshmi S of the Regional Cancer Centre, Trivandrum, for the FACS Facility. AST, HV and SK acknowledge the facilities provided by SCTIMST.

## Author contributions

AST, HV: performed experiments, collected and analysed data and were involved in the preparation of the manuscript. AST and SK: developed the concept, designed experiments, analysed data and were involved in the preparation of the manuscript.

## Supplementary Information

**Supplementary Figure S1** https://doi.org/10.6084/m9.figshare.11371227.v1

## References

1. Anupama V, George M, Dhanesh SB, Chandran A, James J, Shivakumar K. Molecular mechanisms in H_2_O_2_-induced increase in AT1 receptor gene expression in cardiac fibroblasts: A role for endogenously generated Angiotensin II. J Mol Cell Cardiol 97: 295–305, 2016.

2. Booz GW, Baker KM. Actions of Angiotensin II on Isolated Cardiac Myocytes. Heart Fail Rev 3: 125–130, 1998.

3. Bouzegrhane F, Thibault G. Is angiotensin II a proliferative factor of cardiac fibroblasts? Cardiovasc Res 53: 304–312, 2002.

4. Chai J, Norng M, Tarnawski AS, Chow J. A critical role of serum response factor in myofibroblast differentiation during experimental oesophageal ulcer healing in rats. Gut 56: 621–630, 2007.

5. Chen Q, Xie W, Kuhn DJ, Voorhees PM, Lopez-Girona A, Mendy D, Corral LG, Krenitsky VP, Xu W, Moutouh-de Parseval L, Webb DR, Mercurio F, Nakayama KI, Nakayama K, Orlowski RZ. Targeting the p27 E3 ligase SCF(Skp2) results in p27- and Skp2-mediated cell-cycle arrest and activation of autophagy. Blood 111: 4690–4699, 2008.

6. Cowling RT, Yeo SJ, Kim IJ, Park JI, Gu Y, Dalton ND, Peterson KL, Greenberg BH. Discoidin domain receptor 2 germline gene deletion leads to altered heart structure and function in the mouse. Am J Physiol Heart Circ Physiol 307: H773–781, 2014.

7. Crider BJ, Risinger GM, Haaksma CJ, Howard EW, Tomasek JJ. Myocardin-related transcription factors A and B are key regulators of TGF-β1-induced fibroblast to myofibroblast differentiation. J Invest Dermatol 131: 2378–2385, 2011.

8. Duncan JS, Whittle MC, Nakamura K, Abell AN, Midland AA, Zawistowski JS, Johnson NL, Granger DA, Jordan NV, Darr DB, Usary J, Kuan P-F, Smalley DM, Major B, He X, Hoadley KA, Zhou B, Sharpless NE, Perou CM, Kim WY, Gomez SM, Chen X, Jin J, Frye SV, Earp HS, Graves LM, Johnson GL. Dynamic Reprogramming of the Kinome in Response to Targeted MEK Inhibition in Triple-Negative Breast Cancer. Cell 149: 307–321, 2012.

9. Esnault C, Gualdrini F, Horswell S, Kelly G, Stewart A, East P, Matthews N, Treisman R. ERK-Induced Activation of TCF Family of SRF Cofactors Initiates a Chromatin Modification Cascade Associated with Transcription. Mol Cell 65: 1081–1095.e5, 2017.

10. Fan D, Takawale A, Lee J, Kassiri Z. Cardiac fibroblasts, fibrosis and extracellular matrix remodeling in heart disease. Fibrogenesis Tissue Repair 5: 15, 2012.

11. Frescas D, Pagano M. Deregulated proteolysis by the F-box proteins SKP2 and beta-TrCP: tipping the scales of cancer. Nat Rev Cancer 8: 438–449, 2008.

12. Furtado MB, Nim HT, Boyd SE, Rosenthal NA. View from the heart: cardiac fibroblasts in development, scarring and regeneration. Development 143: 387–397, 2016.

13. Gao Y, Chu M, Hong J, Shang J, Xu D. Hypoxia induces cardiac fibroblast proliferation and phenotypic switch: a role for caveolae and caveolin-1/PTEN mediated pathway. Journal of Thoracic Disease 6: 1458–1468, 2014.

14. George M, Vijayakumar A, Dhanesh SB, James J, Shivakumar K. Molecular basis and functional significance of Angiotensin II-induced increase in Discoidin Domain Receptor 2 gene expression in cardiac fibroblasts. J Mol Cell Cardiol 90: 59–69, 2016.

15. Gouldsborough I, Lindop GBM, Ashton N. Renal renin-angiotensin system activity in naturally reared and cross-fostered spontaneously hypertensive rats. Am J Hypertens 16: 864–869, 2003.

16. Grither WR, Longmore GD. Inhibition of tumor–microenvironment interaction and tumor invasion by small-molecule allosteric inhibitor of DDR2 extracellular domain. PNAS 115: E7786–E7794, 2018.

17. Hammerman PS, Sos ML, Ramos AH, Xu C, Dutt A, Zhou W, Brace LE, Woods BA, Lin W, Zhang J, Deng X, Lim SM, Heynck S, Peifer M, Simard JR, Lawrence MS, Onofrio RC, Salvesen HB, Seidel D, Zander T, Heuckmann JM, Soltermann A, Moch H, Koker M, Leenders F, Gabler F, Querings S, Ansén S, Brambilla E, Brambilla C, Lorimier P, Brustugun OT, Helland A, Petersen I, Clement JH, Groen H, Timens W, Sietsma H, Stoelben E, Wolf J, Beer DG, Tsao MS, Hanna M, Hatton C, Eck MJ, Janne PA, Johnson BE, Winckler W, Greulich H, Bass AJ, Cho J, Rauh D, Gray NS, Wong K-K, Haura EB, Thomas RK, Meyerson M. Mutations in the DDR2 kinase gene identify a novel therapeutic target in squamous cell lung cancer. Cancer Discov 1: 78–89, 2011.

18. Hanatani A, Yoshiyama M, Kim S, Omura T, Toda I, Akioka K, Teragaki M, Takeuchi K, Iwao H, Takeda T. Inhibition by angiotensin II type 1 receptor antagonist of cardiac phenotypic modulation after myocardial infarction. Journal of Molecular and Cellular Cardiology 27: 1905–1914, 1995.

19. Hautmann Martina B., Thompson Maria M., Swartz Ellen A., Olson Eric N., Owens Gary K. Angiotensin II–Induced Stimulation of Smooth Muscle α-Actin Expression by Serum Response Factor and the Homeodomain Transcription Factor MHox. Circulation Research 81: 600–610, 1997.

20. Hayashi K, Watanabe B, Nakagawa Y, Minami S, Morita T. RPEL proteins are the molecular targets for CCG-1423, an inhibitor of Rho signaling. PLoS ONE 9: e89016, 2014.

21. Henley SA, Dick FA. The retinoblastoma family of proteins and their regulatory functions in the mammalian cell division cycle. Cell Div 7: 10, 2012.

22. Humeres C, Frangogiannis NG. Fibroblasts in the Infarcted, Remodeling, and Failing Heart. J Am Coll Cardiol Basic Trans Science 4: 449–467, 2019.

23. Jia S, Agarwal M, Yang J, Horowitz JC, White ES, Kim KK. Discoidin Domain Receptor 2 Signaling Regulates Fibroblast Apoptosis through PDK1/Akt. Am J Respir Cell Mol Biol 59: 295–305, 2018

24. Kajstura J, Cigola E, Malhotra A, Li P, Cheng W, Meggs LG, Anversa P. Angiotensin II induces apoptosis of adult ventricular myocytes in vitro. J Mol Cell Cardiol 29: 859–870, 1997.

25. Kawai I, Hisaki T, Sugiura K, Naito K, Kano K. Discoidin domain receptor 2 (DDR2) regulates proliferation of endochondral cells in mice. Biochem Biophys Res Commun 427: 611–617, 2012.

26. Khattar E, Kumar V. Mitogenic regulation of p27(Kip1) gene is mediated by AP-1 transcription factors. J Biol Chem 285: 4554–4561, 2010.

27. Kumaran C, Shivakumar K. Calcium- and superoxide anion-mediated mitogenic action of substance P on cardiac fibroblasts. Am J Physiol Heart Circ Physiol 282: H1855–1862, 2002.

28. Labrador JP, Azcoitia V, Tuckermann J, Lin C, Olaso E, Manes S, Bruckner K, Goergen JL, Lemke G, Yancopoulos G, Angel P, Martínez C, Klein R. The collagen receptor DDR2 regulates proliferation and its elimination leads to dwarfism. EMBO Rep 2: 446–452, 2001.

29. Lees SJ, Childs TE, Booth FW. Age-dependent FOXO regulation of p27Kip1 expression via a conserved binding motif in rat muscle precursor cells. American Journal of Physiology-Cell Physiology 295: C1238–C1246, 2008.

30. Leitinger B. Discoidin domain receptor functions in physiological and pathological conditions. Int Rev Cell Mol Biol 310: 39–87, 2014.

31. Li L, Yue Z, Wan X, Zhang G, Song S, Bai X, Jiao Y, Ju Y, Li J. Alteration of discoidin domain receptor-2 expression: possible role in peroxynitrite-induced apoptosis in human cerebral vascular smooth muscle cells. Mol Cell Toxicol 8: 401–406, 2012.

32. Lin K-L, Chou C-H, Hsieh S-C, Hwa S-Y, Lee M-T, Wang F-F. Transcriptional upregulation of DDR2 by ATF4 facilitates osteoblastic differentiation through p38 MAPK-mediated Runx2 activation. Journal of Bone and Mineral Research 25: 2489–2503, 2010.

33. Luo Z, Chen S, Gill P, Kawada N, Andresen BT, Welch WJ, Jose PA, Wilcox CS. Angiotensin II induced reactive oxygen species modulate smooth muscle α-actin gene expression via increased SRF binding to CArG cis elements in SHR renal microvascular smooth muscle cells. The FASEB Journal 20: A1148–A1148, 2006.

34. Luo Z, Liu H, Sun X, Guo R, Cui R, Ma X, Yan M. RNA Interference against Discoidin Domain Receptor 2 Ameliorates Alcoholic Liver Disease in Rats. PLOS ONE 8: e55860, 2013.

35. Lv T, Du Y, Cao N, Zhang S, Gong Y, Bai Y, Wang W, Liu H. Proliferation in cardiac fibroblasts induced by β 1 -adrenoceptor autoantibody and the underlying mechanisms. Sci Rep 6: 1–15, 2016.

36. Maeyama M, Koga H, Selvendiran K, Yanagimoto C, Hanada S, Taniguchi E, Kawaguchi T, Harada M, Ueno T, Sata M. Switching in discoid domain receptor expressions in SLUG-induced epithelial-mesenchymal transition. Cancer 113: 2823–2831, 2008.

37. Malumbres M, Barbacid M. Cell cycle, CDKs and cancer: a changing paradigm. Nat Rev Cancer 9: 153–166, 2009.

38. Mayorga M, Bahi N, Ballester M, Comella JX, Sanchis D. Bcl-2 Is a Key Factor for Cardiac Fibroblast Resistance to Programmed Cell Death. J Biol Chem 279: 34882–34889, 2004.

39. Obaya AJ, Sedivy JM. Regulation of cyclin-Cdk activity in mammalian cells. Cellular and Molecular Life Sciences (CMLS) 59: 126–142, 2002.

40. Oishi Y, Ozono R, Yoshizumi M, Akishita M, Horiuchi M, Oshima T. AT2 receptor mediates the cardioprotective effects of AT1 receptor antagonist in post-myocardial infarction remodeling. Life Sciences 80: 82–88, 2006.

41. Olaso E, Arteta B, Benedicto A, Crende O, Friedman SL. Loss of Discoidin Domain Receptor 2 Promotes Hepatic Fibrosis after Chronic Carbon Tetrachloride through Altered Paracrine Interactions between Hepatic Stellate Cells and Liver-Associated Macrophages. The American Journal of Pathology 179: 2894–2904, 2011.

42. Olaso E, Ikeda K, Eng FJ, Xu L, Wang LH, Lin HC, Friedman SL. DDR2 receptor promotes MMP-2-mediated proliferation and invasion by hepatic stellate cells. J Clin Invest 108: 1369–1378, 2001.

43. Olaso E, Labrador J-P, Wang L, Ikeda K, Eng FJ, Klein R, Lovett DH, Lin HC, Friedman SL. Discoidin domain receptor 2 regulates fibroblast proliferation and migration through the extracellular matrix in association with transcriptional activation of matrix metalloproteinase-2. J Biol Chem 277: 3606–3613, 2002.

44. Olsen MB, Hildrestrand GA, Scheffler K, Vinge LE, Alfsnes K, Palibrk V, Wang J, Neurauter CG, Luna L, Johansen J, Øgaard JDS, Ohm IK, Slupphaug G, Kuśnierczyk A, Fiane AE, Brorson S-H, Zhang L, Gullestad L, Louch WE, Iversen PO, Østlie I, Klungland A, Christensen G, Sjaastad I, Satrom P, Yndestad A, Aukrust P, Bjørås M, Finsen AV. NEIL3-Dependent Regulation of Cardiac Fibroblast Proliferation Prevents Myocardial Rupture. Cell Reports 18: 82–92, 2017.

45. Olson EN, Nordheim A. Linking actin dynamics and gene transcription to drive cellular motile functions. Nat Rev Mol Cell Biol 11: 353–365, 2010.

46. Olson ER, Naugle JE, Zhang X, Bomser JA, Meszaros JG. Inhibition of cardiac fibroblast proliferation and myofibroblast differentiation by resveratrol. American Journal of Physiology-Heart and Circulatory Physiology 288: H1131–H1138, 2005.

47. Parmacek Michael S. Myocardin-Related Transcription Factors. Circulation Research 100: 633–644, 2007.

48. Philip L, Shivakumar K. cIAP-2 protects cardiac fibroblasts from oxidative damage: an obligate regulatory role for ERK1/2 MAPK and NF-κB. J Mol Cell Cardiol 62: 217–226, 2013.

49. Pramod S, Shivakumar K. Mechanisms in cardiac fibroblast growth: an obligate role for Skp2 and FOXO3a in ERK1/2 MAPK-dependent regulation of p27kip1. Am J Physiol Heart Circ Physiol 306: H844–855, 2014.

50. Prescott MF, Webb RL, Reidy MA. Angiotensin-converting enzyme inhibitor versus angiotensin II, AT1 receptor antagonist. Effects on smooth muscle cell migration and proliferation after balloon catheter injury. Am J Pathol 139: 1291–1296, 1991.

51. Santo EE, Stroeken P, Sluis PV, Koster J, Versteeg R, Westerhout EM. FOXO3a is a major target of inactivation by PI3K/AKT signaling in aggressive neuroblastoma. Cancer Res 73: 2189–2198, 2013.

52. Schratt G, Philippar U, Hockemeyer D, Schwarz H, Alberti S, Nordheim A. SRF regulates Bcl-2 expression and promotes cell survival during murine embryonic development. EMBO J 23: 1834–1844, 2004.

53. Sladek T, Sladkova J, Kolar F, Papousek F, Cicutti N, Korecky B, Rakusan K. The effect of AT1 receptor antagonist on chronic cardiac response to coronary artery ligation in rats. Cardiovasc Res 31: 568–576, 1996.

54. Small EM. The actin-MRTF-SRF gene regulatory axis and myofibroblast differentiation. J Cardiovasc Transl Res 5: 794–804, 2012.

55. Sopel MJ, Rosin NL, Lee TDG, Legare J-F. Myocardial fibrosis in response to Angiotensin II is preceded by the recruitment of mesenchymal progenitor cells. Lab Invest 91: 565–578, 2011.

56. Taurin S, Sandbo N, Yau DM, Sethakorn N, Kach J, Dulin NO. Phosphorylation of Myocardin Extracellular Signal-regulated Kinase. J Biol Chem 284: 33789–33794, 2009.

57. Thannickal VJ, Horowitz JC. Evolving Concepts of Apoptosis in Idiopathic Pulmonary Fibrosis. Proc Am Thorac Soc 3: 350–356, 2006.

58. Ushakumary MG, Wang M, V H, Titus AS, Zhang J, Liu L, Monticone R, Wang Y, Mattison JA, Cabo R de, Lakatta EG, Kailasam S. Discoidin domain Receptor 2: A determinant of metabolic syndrome-associated arterial fibrosis in non-human primates. PLOS ONE 14: e0225911, 2019.

59. V H, Titus AS, Cowling RT, Kailasam S. Collagen receptor cross-talk determines α-smooth muscle actin-dependent collagen gene expression in angiotensin II-stimulated cardiac fibroblasts. J Biol Chem 294: 19723–19739, 2019.

60. Valiathan RR, Marco M, Leitinger B, Kleer CG, Fridman R. DISCOIDIN DOMAIN RECEPTOR TYROSINE KINASES: NEW PLAYERS IN CANCER PROGRESSION. Cancer Metastasis Rev 31: 295–321, 2012.

61. Wall SJ, Werner E, Werb Z, DeClerck YA. Discoidin Domain Receptor 2 Mediates Tumor Cell Cycle Arrest Induced by Fibrillar Collagen. J Biol Chem 280: 40187–40194, 2005.

62. Wang D-Z, Li S, Hockemeyer D, Sutherland L, Wang Z, Schratt G, Richardson JA, Nordheim Olson EN. Potentiation of serum response factor activity by a family of myocardin-related transcription factors. Proc Natl Acad Sci USA 99: 14855–14860, 2002.

63. Weidinger C, Krause K, Klagge A, Karger S, Fuhrer D. Forkhead box-O transcription factor: critical conductors of cancer’s fate. Endocr Relat Cancer 15: 917–929, 2008.

64. Werth D, Grassi G, Konjer N, Dapas B, Farra R, Giansante C, Kandolf R, Guarnieri G, Nordheim A, Heidenreich O. Proliferation of human primary vascular smooth muscle cells depends on serum response factor. European Journal of Cell Biology 89: 216–224, 2010.

65. Yang J-Y, Hung M-C. A new fork for clinical application: targeting forkhead transcription factors in cancer. Clin Cancer Res 15: 752–757, 2009.

66. Zhang S, Huan W, Wei H, Shi J, Fan J, Zhao J, Shen A, Teng H. FOXO3a/p27kip1 expression and essential role after acute spinal cord injury in adult rat. J Cell Biochem 114: 354–365, 2013.

67. Zhou N, Lee J-J, Stoll S, Ma B, Wiener R, Wang C, Costa KD, Qiu H. Inhibition of SRF/myocardin reduces aortic stiffness by targeting vascular smooth muscle cell stiffening in hypertension. Cardiovasc Res 113: 171–182, 2017.

