## Supplementary figures and images for "Co-ordinated regulation of cell survival and cell cycle pathways by DDR2-dependent SRF transcription factor in cardiac fibroblasts" (from another version of this preprint: version 1, 2019-11-27)

### Supplementary Figure 2

**Supplementary Figure S2: Representative scatter plot images of Annexin-PI by flow cytometry**

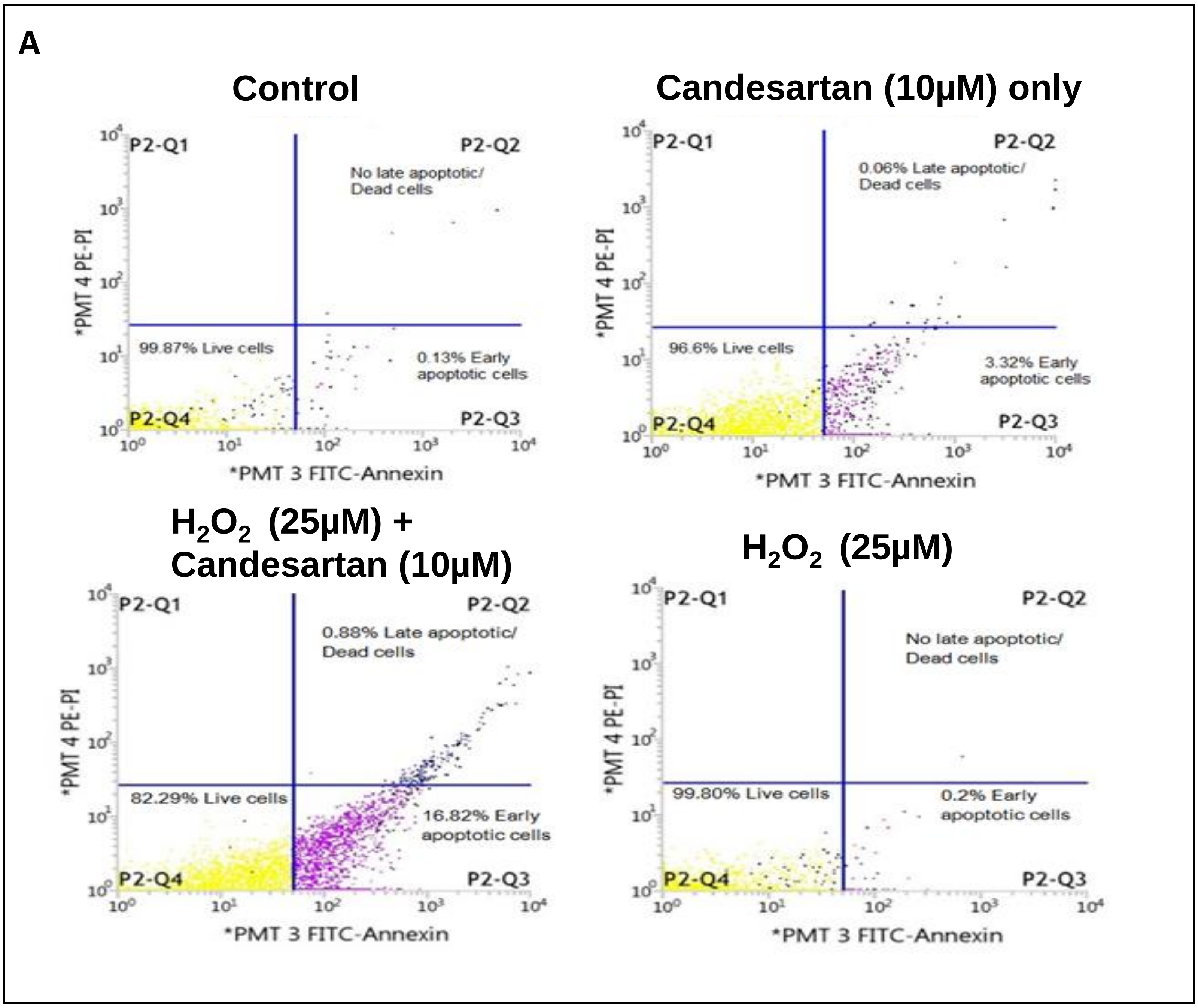


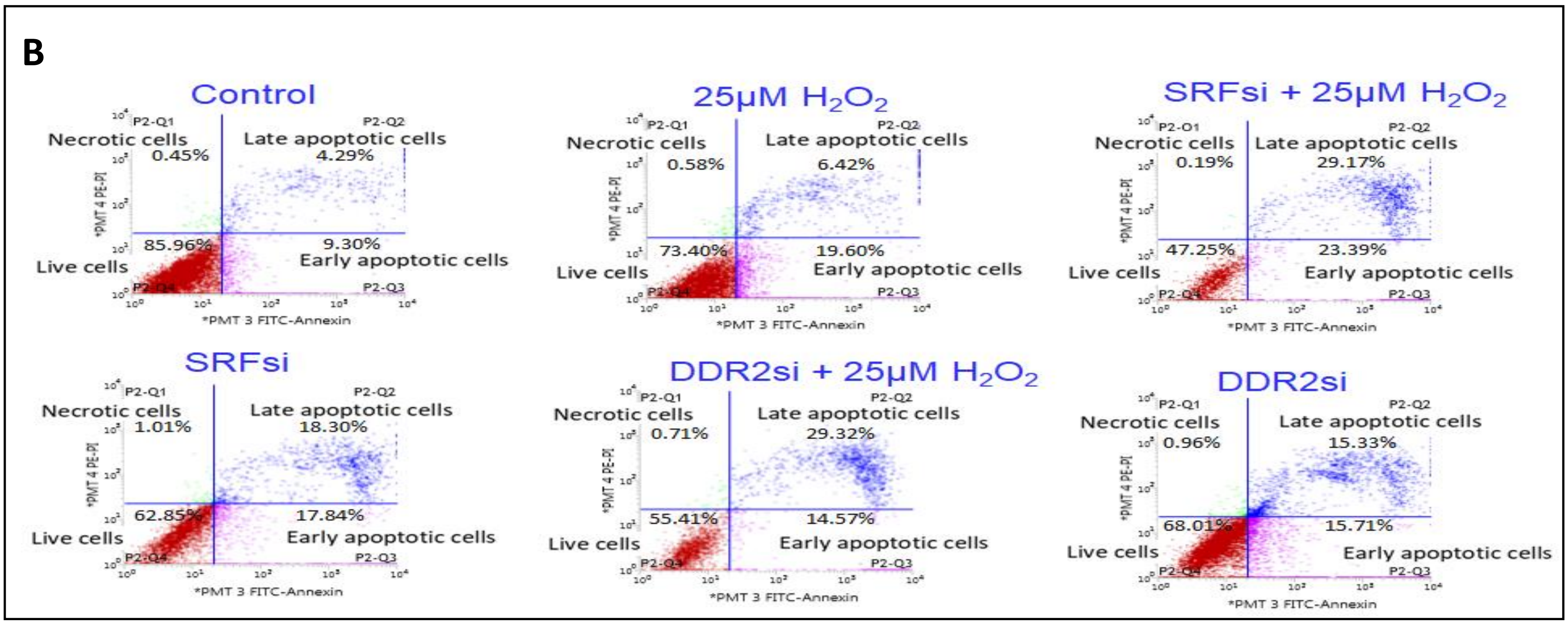
